# A spatial topology atlas of type I myofibers in porcine skeletal muscle reveals local slow-oxidative niches associated with meat quality traits

**DOI:** 10.64898/2026.10.08.757308

**Authors:** Zhaoli Shen, Yuehao Jiao, Geng Zhang, Jin Chai, Guoliang Han, Chenggang Gao, Mingzhou Li, Jin Long

## Abstract

Skeletal muscle metabolism and meat quality are usually interpreted through myofiber type composition, fiber size, and histochemical state. Whether the spatial topology of myofibers constitutes an independent phenotype layer remains poorly quantified. Here, using succinate dehydrogenase (SDH)-stained cross-sectional images from multiple pig breeds and more than 40 skeletal muscle sites, we established a multiscale spatial phenotyping framework for type I myofibers. To enable scalable instance-level quantification, we developed an improved Mask R-CNN segmentation workflow, trained with SAM-assisted and manually corrected annotations, and optimized for dense, small, irregular, and variably stained muscle fibers. This framework captures singlefiber morphology and staining, local contact, connected-cluster architecture, diffusion-ring neighborhoods, and global tissue configuration. Type I myofiber abundance was separable from spatial organization capacity: some muscles formed stronger-than-expected local connectivity and clustering despite modest type I fiber abundance. Spatial transcriptomic integration further showed that slow-oxidative genes and functional modules were nonrandomly organized and formed local high-expression domains in type I fiber-enriched, high-contact neighborhoods. Across muscle sites and breeds, skeletal muscles displayed distinguishable type I fiber spatial topology identities, which were remodeled by breed background in a muscle-specific manner. In samples with matched meat-quality measurements, spatial topology features improved the explanation of pH, meat color, conductivity, and intramuscular fat. These results suggest that skeletal muscle should be modeled as a spatially organized tissue system rather than a simple mixture of fiber types.

**Highlights:**

- A multiscale spatial phenotyping framework was established for porcine type I myofibers.
- Type I fiber topology organization are separable from abundance and affects muscle fiber amplification and gene expression.
- Breed background remodels type I fiber spatial topology in a muscle-specific manner.
- Spatial topology features improve explanation of pH, meat color, conductivity, and IMF
- The improved deep learning algorithm supports more efficient muscle fiber segmentation and phenotype extraction.

## Introduction

Livestock skeletal muscle is the primary component of meat products, and its development, fiber-type composition, metabolic state, and tissue architecture jointly determine meat yield and quality-related traits [1-5]. Systematic characterization of livestock skeletal muscle is therefore essential for understanding the biological basis of meat-production traits and for supporting efficient and sustainable production [6-11]. In parallel, porcine skeletal muscle resembles human muscle in structural and biochemical properties, making pigs a valuable model for muscle biology, atrophy, metabolic disorders, and related diseases [12,13].

Myofibers are the basic structural and functional units of skeletal muscle. Their type composition, size, and density reflect developmental status, contractile properties, and metabolic function [14-18]. Previous population-level studies in pigs have linked cross-sectional myofiber phenotypes, including fiber-type proportion, crosssectional area, and density, to fat deposition, meat color, water-holding capacity, postmortem pH, and sensory quality [19-25]; related reviews are available in [8,19,26]. Cross-species studies further indicate that different myofiber types are not randomly arranged, but instead show clustering, stratification, or neighborhood-specific organization in birds, cattle, pigs, mice, and humans [8,13,17-18]. These patterns vary with anatomical site, development, aging, and neural activity [28-39]. Spatial myofiber arrangement may therefore be linked to local cell-cell interactions, gene-expression states, and tissue-level traits [28-39].

However, existing phenotyping has focused largely on fiber-type proportion and cross-sectional area. Studies of myofiber spatial distribution have remained relatively limited, often stopping at the question of whether fiber types are clustered or randomly distributed, without a multiscale framework spanning single fibers, local microenvironments, and global tissue configuration [7-8,11,14,16,40-48]. In addition, myofiber phenotype extraction remains constrained by image-analysis performance. Classical computer-vision methods are sensitive to noise and staining variation [49-60]. Although deep learning has accelerated cellular instance segmentation [40-45,47-48,50,61-84], skeletal muscle images contain large numbers of densely packed and relatively small targets, posing continuing challenges for segmentation accuracy, computational efficiency, and downstream phenotype quantification.

Here, we developed a multiscale spatial phenotyping framework for skeletal muscle cross-sections. The framework extends conventional measurements of fiber proportion and size to quantitative representations of single-fiber morphology, local neighborhood structure, and whole-tissue spatial configuration. We applied this framework to histological images from multiple pig breeds and skeletal muscle sites to assess the separability of type I fiber abundance and spatial topology, to compare spatial organization across muscles and breed backgrounds, and to link myofiber topology with local molecular state and tissue-level traits using spatial transcriptomics and meat-quality measurements. To enable scalable application, we also optimized dense myofiber instance segmentation, establishing an integrated analysis path from tissue images to spatial phenotypes, molecular spatial states, and tissue-level outputs.

We show that type I myofiber spatial topology represents an independent tissue-structural layer beyond fiber abundance and size. This topology is linked to local slow-oxidative molecular programs, varies across muscle sites and breed backgrounds, and contributes to meat-quality traits. The proposed deep learning segmentation workflow improves scalable conversion of histological images into reproducible multiscale muscle phenotypes.

## Results

### 1. Establishing a multiscale spatial phenotyping framework for type I myofibers

To systematically quantify the spatial organization of type I myofibers in skeletal muscle, we first established a multiscale phenotyping framework that extends conventional measurements of fiber proportion and mean size to an integrated representation of single-fiber attributes, local microenvironments, and global tissue configuration (Figure 1, Table S1). The main data were SDH-stained cross-sectional images from multiple pig breeds (Table S2) and skeletal muscle sites (Table S3). After image-quality filtering, these images formed a first dataset for multi-site phenotypic comparison and a second dataset with matched tissue-level trait measurements (Table S4).

**Figure 1.**
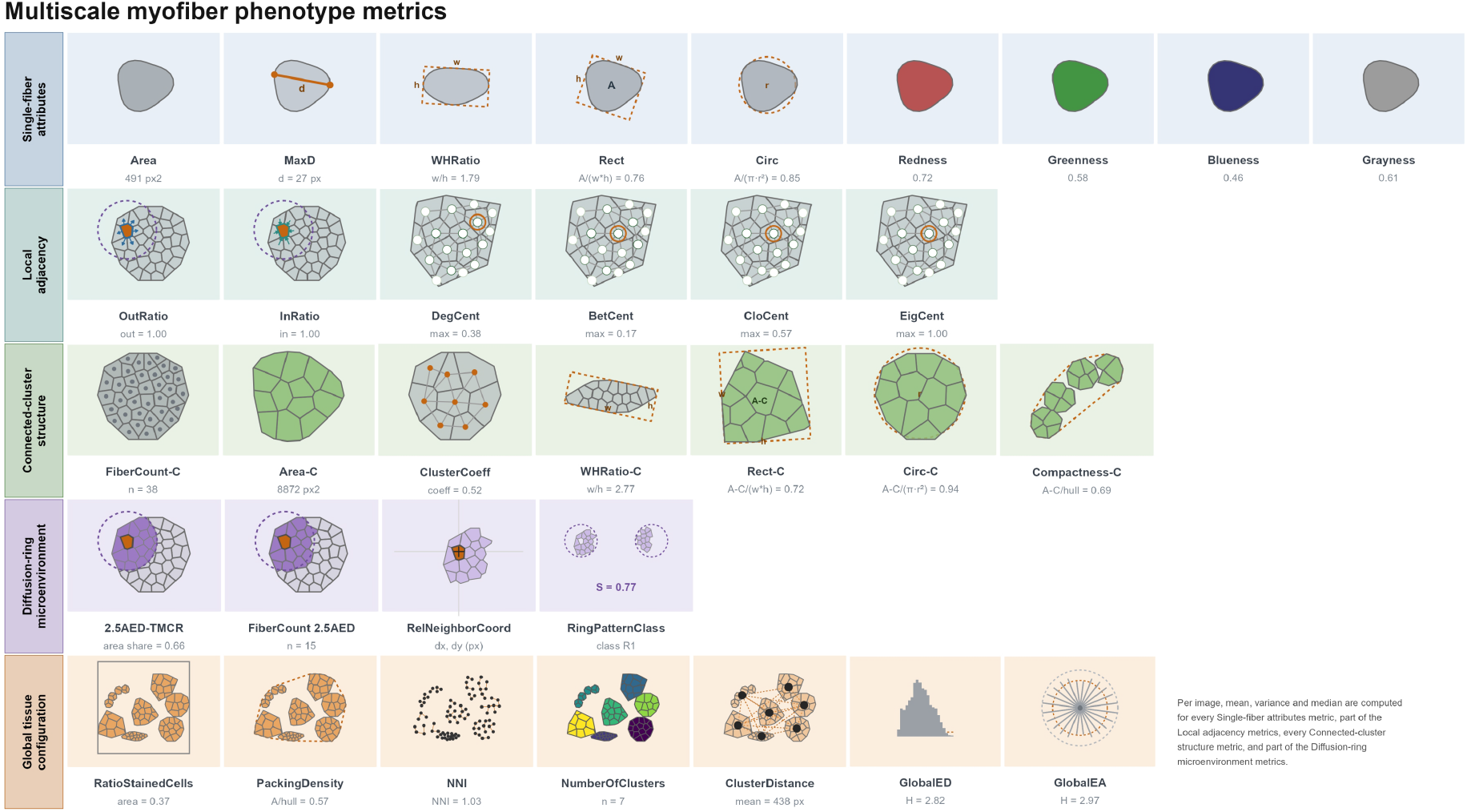
Multiscale spatial phenotyping framework for type I myofibers. The framework converts tissue images into multilevel descriptors of single fibers, local neighborhoods, connected clusters, and global tissue configuration.

To obtain reliable objects for phenotype calculation, we first used Segment Anything for preliminary myofiber-contour segmentation and then manually corrected the contours in Labelme, yielding high-confidence annotations of type I myofibers (Figure S1A,B). Unless otherwise stated, “myofiber” below refers to SDH-positive type I myofibers. Based on these contours, each tissue image was converted into a computable collection of spatial objects for downstream multiscale feature extraction.

At the single-fiber scale, the framework describes both intrinsic fiber attributes and local topological position. Intrinsic attributes include area, diameter, aspect ratio, circularity, rectangularity, orientation, and staining intensity, which capture morphology and metabolic staining of individual fibers. Topological attributes are derived from fiber-fiber adjacency and quantify how strongly each fiber contacts neighboring fibers of the same type and where it sits within the local network. Thus, an individual myofiber is no longer treated as an isolated geometric object, but as an object embedded in its local tissue context.

At the local microenvironment scale, we used two complementary neighborhood definitions. The first is based on direct adjacency between fiber contours, from which connected clusters of same-type myofibers are constructed and quantified by cluster size, shape, compactness, clustering coefficient, and graph centrality. The second defines a diffusion-ring neighborhood centered on each myofiber and measures same-type fiber area fraction, local enrichment, and spatial displacement within the circular neighborhood. The former emphasizes direct contact and clustering; the latter describes same-type fiber enrichment and distribution in a local spatial field. The consistent clustering patterns (Figure S1) observed under both definitions suggest that type I myofiber microenvironments are not randomly assembled, but may represent intrinsically organized local tissue units with potential structural and functional relevance.

At the whole-tissue scale, single-fiber and local-neighborhood features are summarized into image-level spatial phenotypes. In addition to distributional statistics such as mean, variance, and median, we calculated nearest-neighbor index, stained-fiber area fraction, connected-cluster number, inter-cluster distance, packing density, and entropy derived from global distance and angle matrices. These features describe the structural state of the entire skeletal muscle cross-section from the perspectives of abundance, density, aggregation, tissue complexity, and spatial heterogeneity.

Through this framework, each tissue image is represented as a multiscale phenotype matrix spanning fiber abundance, geometry, and spatial topology. This matrix provides a unified quantitative basis for testing whether type I myofiber spatial topology is independent of fiber proportion and size, and for linking local molecular state, muscle-site differences, breed background, and tissue-level traits. Figure S2 presents a simple example to illustrate that although the proportions of type I fibers are similar, their spatial distributions are not consistent.

### 2. Local topology of type I myofibers reflects phenotype autocorrelation and neighbor-dependent cluster expansion

After establishing the phenotyping framework, we first examined whether type I myofiber topology reflects interpretable tissue states rather than merely geometric consequences of fiber number. Using SDH images from multiple skeletal muscles of Large White (LW) and Rongchang (RC) pigs, we analyzed the relationships among type I myofiber abundance, global distribution, local adjacency, and connected-cluster structure.

At the global distribution level, type I fiber number was positively associated with nearest-neighbor index and spatial packing density, but negatively associated with connected-cluster number and inter-cluster distance (Figure 2A). This pattern suggests that, as type I fiber number increases, newly added fibers are more likely to join or expand existing local structures than to form uniformly distributed isolated clusters. At the local scale, increased type I fiber proportion was accompanied by stronger same-type contact, reflected by higher InRatio, OutRatio, 2.5AED-TMCR, within-cluster fiber count, and clustering coefficient (Figure 2A,B). These results support a neighbor-dependent clustering expansion pattern in which increased type I fiber abundance is coupled to enhanced local contact and same-type enrichment.

**Figure 2.**
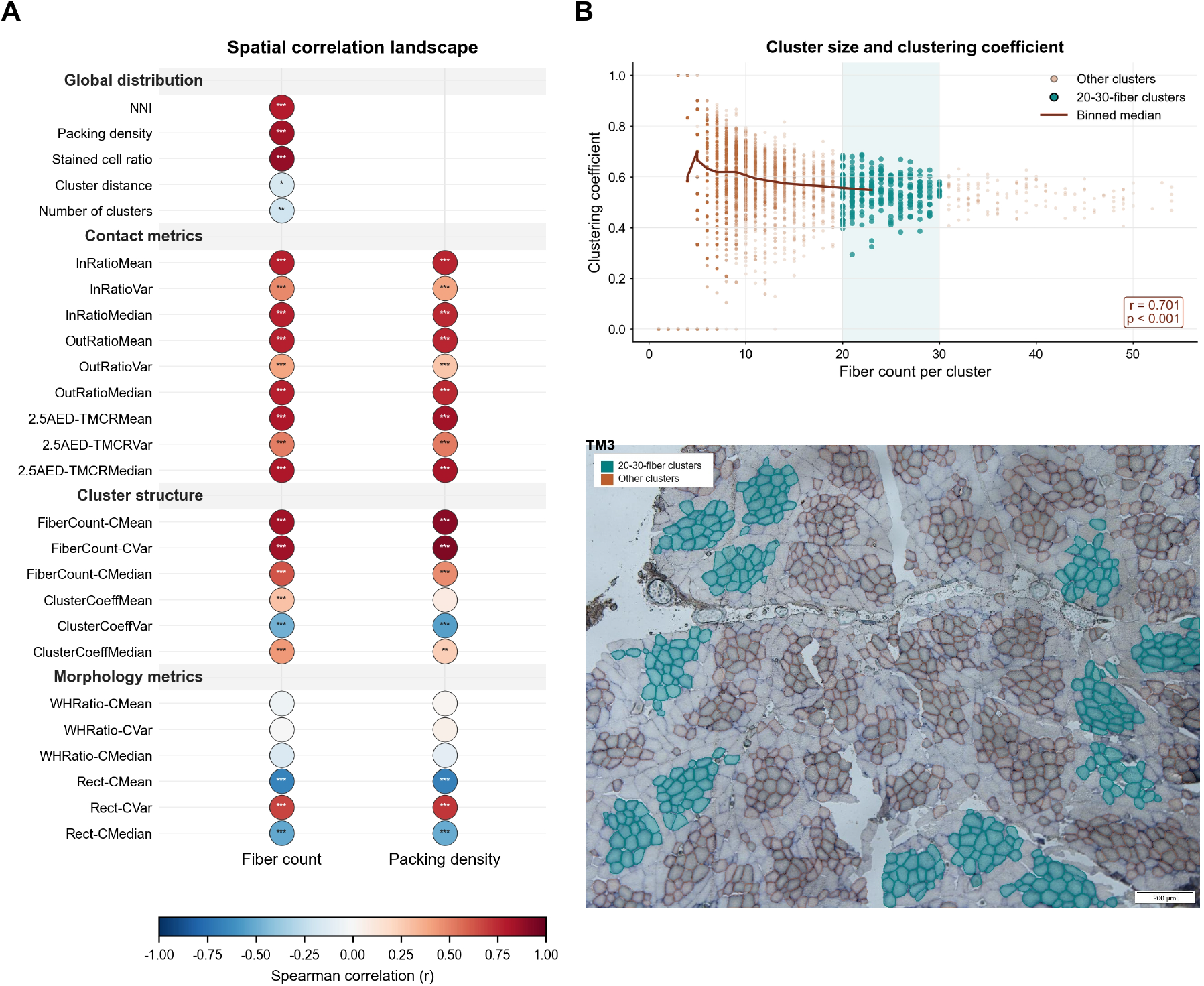
Spatial relationships among type I myofiber abundance, local adjacency, and connected-cluster structure. **(A)** Integrated correlation landscape showing associations between type I myofiber abundance- related traits and spatial metrics, including global distribution metrics, local contact degree metrics, connected-cluster fiber count, clustering coefficient, and morphology-related metrics. Circle color indicates Spearman correlation direction and magnitude, circle size indicates absolute correlation strength, and asterisks indicate statistical significance. **(B)** Relationship between intra-cluster fiber count and clustering coefficient across connected clusters. Clusters containing 20-30 fibers are highlighted, and the representative TM3 tissue image shows the spatial distribution of 20-30-fiber connected clusters relative to other connected clusters.

This neighbor-dependent expansion was not a simple linear accumulation. As the number of fibers in a connected cluster increased, the within-cluster clustering coefficient generally increased, but the growth appeared constrained by cluster size, local spatial capacity, and tissue geometry (Figure 2B). Thus, type I myofiber spatial organization is more likely achieved through the formation and expansion of multiple local connected units than through unlimited enlargement of a single large cluster. This observation provides a basis for distinguishing increased fiber abundance from enhanced topological organization.

At the single-fiber scale, local topological position was coupled to the phenotype of type I myofibers themselves. Site-specific analyses in the longissimus dorsi and psoas major further supported associations between Area, muscle fiber count, Packing Density, and morphology, cluster structure, and contact metrics. (Figure S3). Together, these results suggest that type I fibers located in high-contact or high-centrality microenvironments tend to exhibit stronger local phenotypic features.

### 3. Type I fiber spatial topology provides an independent explanatory layer for tissue-level traits

The preceding results show that type I fiber spatial topology has molecular meaning and is remodeled in muscle- and breed-specific manners. To test whether these microstructural features connect to tissue-level outputs, we analyzed a second set of SDH images with matched trait records. This analysis included only samples with valid tissue images, extracted type I fiber phenotypes, and trait measurements, mainly LDM and PMM. Images without matched trait records were excluded from PLS and correlation analyses.

We associated multiscale type I fiber phenotypes with tissue-level traits including meat color L*, a*, and b*, pH, conductivity at 45-60 min and 24 h, and intramuscular fat (IMF). Partial least squares regression (PLS) was used to evaluate the explanatory capacity of the phenotype matrix for each trait. Variable importance in projection (VIP), Spearman correlation, and control models excluding spatial metrics were used to determine whether topology provides independent information beyond conventional proportion and size metrics.

PLS results showed that many tissue-level traits could be well explained by type I fiber phenotypes, with model R^2^ > 0.5 or mean absolute relative error (MARE) < 20% (Figure 3). The best performance was observed for pH at 24 h, with MARE = 7.10% and R^2^ = 0.85. Because pH at 24 h reflects the terminal postmortem acidification state, this result suggests that type I fiber phenotypes, especially their spatial organization, may be related to local metabolic state and acidification dynamics rather than merely static tissue morphology.

**Figure 3.**
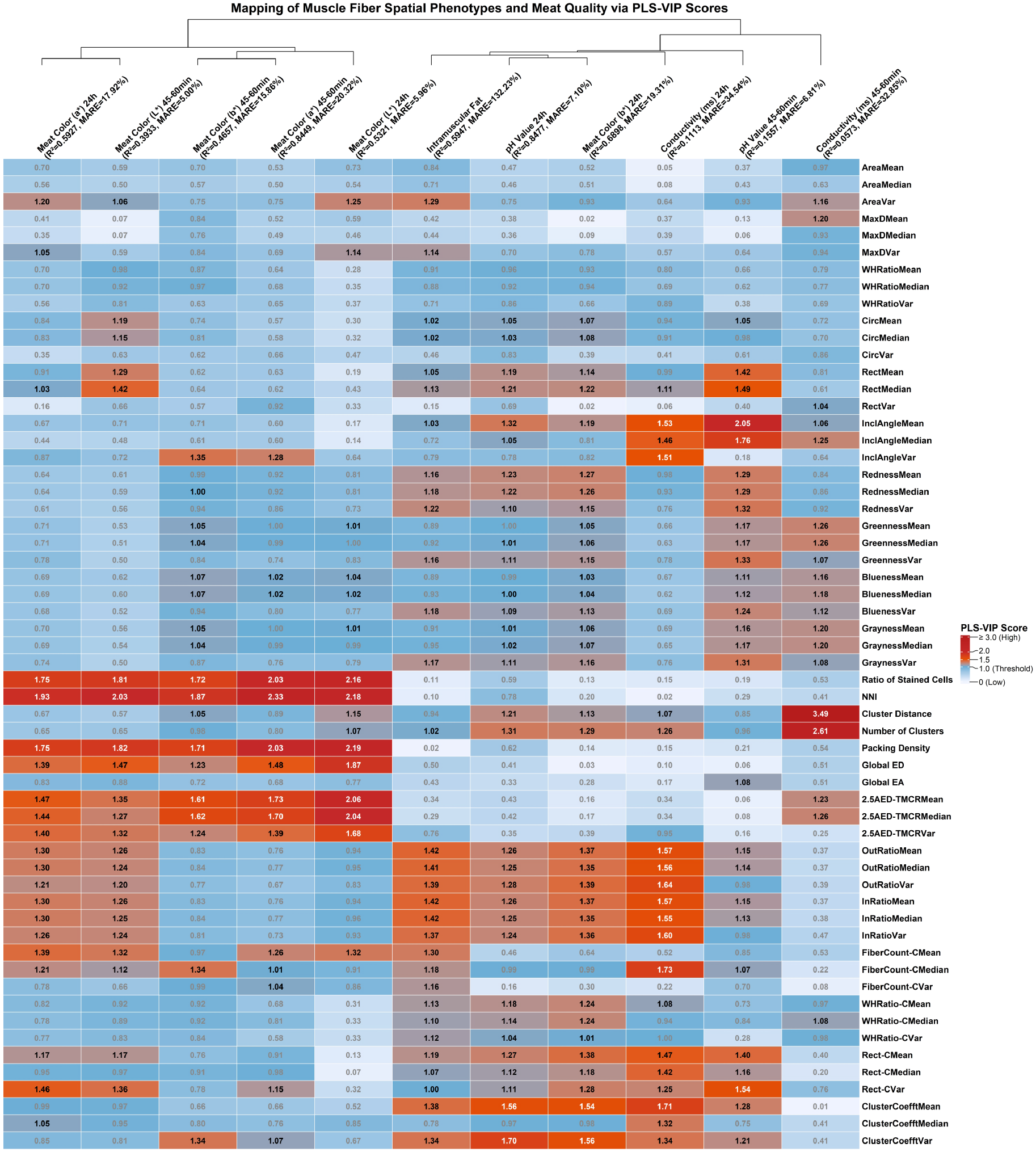
PLS-VIP analysis shows that type I fiber spatial phenotypes explain multiple tissue-level traits.

VIP analysis further showed that connected-cluster architecture, inter-fiber contact, and local enrichment metrics were important explanatory variables for multiple traits (Figure 3). For pH at 24 h, cluster size, within- cluster adjacency, InRatio, OutRatio, and 2.5AED-TMCRMean had high weights. These spatial metrics also contributed strongly to b* at 24 h, IMF, and pH at 45-60 min. By contrast, L*, a*, and early b* values were more closely associated with overall type I fiber proportion, NNI, global compactness and 2.5AED-TMCRMean. The phenotype-trait relationships can therefore be summarized into two patterns: a topology/contact pattern centered on connected clusters and local contact, mainly associated with pH at 24 h, b* at 24 h, and IMF; and an abundance/distribution pattern centered on fiber proportion and global density, mainly associated with color phenotypes such as L* and a*.

Spearman correlation analysis supported these patterns (Figure S4, Figure S5). In general, larger type I fiber clusters, stronger within-cluster adjacency, shorter inter-cluster distances, and higher InRatio/OutRatio were associated with lower pH at 24 h, higher b* at 24 h, and lower IMF. NNI, global compactness, total type I fiber area, and 2.5AED-TMCRMean were positively correlated with a* at 45-60 min/24 h and b* at 45-60 min, but negatively correlated with L* at 45-60 min/24 h. Different spatial metrics therefore do not all point to the same tissue output; rather, they correspond to distinct dimensions including acidification, color, and lipid deposition.

The independent value of spatial metrics became clearer in control models. When spatial distribution, local adjacency, and connected-cluster metrics were removed, leaving only conventional measures such as type I fiber proportion, area, and morphology, the overall performance of PLS models declined (Figure S6). Spatial topology is therefore not a redundant expression of fiber composition. Even if two samples have similar type I fiber proportions or mean areas, whether type I fibers form compact connected clusters, exhibit high local contact, or are enriched within local circular neighborhoods may still influence tissue-level traits.

We also evaluated coarse-grained global spatial-distance and angle metrics derived from Gaussian radial basis function fitting [85]. These metrics showed only weak and unstable correlations with trait differences (Figure S7). This suggests that not all spatial differences have equal explanatory value. Highly globalized spatial distances may mix structural information across levels, whereas connected clusters, local adjacency, and diffusion-ring metrics have clearer histological meaning and are better suited to tracking microstructural sources associated with pH, color, and IMF.

Overall, this section directly links type I fiber spatial topology to tissue-level traits. Section 6 showed that type I fiber abundance and topology capacity are separable; Sections 7 and 8 showed that topology identity is site-specific and remodeled by breed background; and this section shows that these spatial features improve the explanation of pH, meat color, conductivity, and IMF. Skeletal muscle should therefore be understood as a spatially structured system in which type I fiber clustering, adjacency, and local enrichment form an intermediate phenotype layer connecting fiber composition, local molecular state, and tissue output.

### 4. Trait-specific topology scores link spatial organization to bulk transcriptional programs

Following the image-based and spatial-transcriptomic analyses above, we next asked whether topology- derived spatial exposures were associated with transcriptional state in matched bulk RNA-sequencing samples. This analysis was designed as a complementary, phenotype-anchored test of the relationship between tissue organization and molecular programs. We reconstructed seven spatial metrics for 98 exact phenotype-bulk RNA-seq matches spanning 39 muscle sites and applied the prespecified fixed PLS parameters to five reduced spatial inputs: cluster coefficient, out-ratio, cluster distance, fiber count, and a local-density composite score. The composite score integrates directionally harmonized nearest-neighbor index, packing density, and 2.5AEDTMCR information. For each of the 11 meat-quality traits, the full PLS regression coefficients were retained to define a trait-specific topology score (Figure 4A). These scores represent multivariate spatial exposures and should not be interpreted as the independent causal effects of individual metrics.

**Figure 4.**
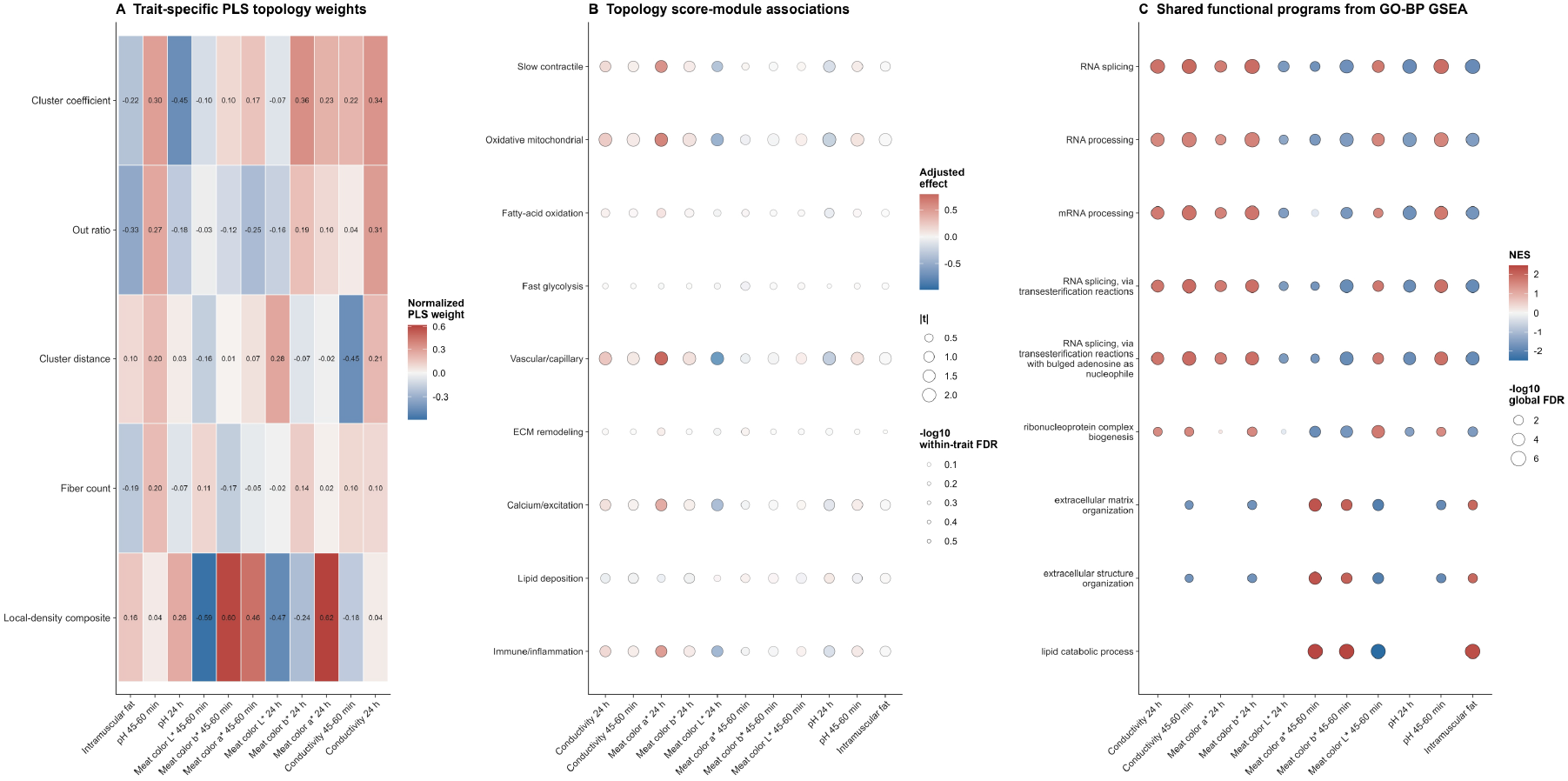
Trait-specific spatial topology scores are associated with bulk transcriptional programs and functional enrichment. **(A)** Trait-specific topology scores derived from fixed PLS regression coefficients. For each of the 11 meat-quality traits, the score was constructed from five reduced spatial inputs: cluster coefficient, out-ratio, cluster distance, fiber count, and the local-density composite score. The local-density composite integrates directionally harmonized nearest-neighbor index, packing density, and 2.5AED-TMCR information. Coefficients were normalized within each trait by the sum of their absolute values. Red and blue indicate positive and negative contributions, respectively.**(B)** Associations between trait-specific topology scores and nine prespecified transcriptional modules. Each point represents one of 99 trait–module combinations estimated using the model module score ∼ 0 + muscle site + topology score + Type I fiber fraction in 98 exact phenotype–bulk RNA-seq matches spanning 39 muscle sites. Point position indicates the adjusted regression effect, point size represents the absolute moderated t statistic, and transparency indicates the strength of the within-trait FDR signal. No association remained significant after trait-wise or global FDR correction.**(C)** Ranked GO Biological Process gene-set enrichment associated with trait-specific topology scores. For each trait, genes were ranked by the moderated t statistic from the primary exact-site model, and GSEA was performed using 10,000 permutations with deterministic trait-specific random seeds. The displayed terms comprise the union of the three highest-ranked terms for each trait, selected by adjusted P value, nominal P value, and absolute normalized enrichment score (NES). Point color indicates NES, point size indicates the negative logarithm of the global FDR across traits and gene sets, and filled versus translucent points indicate global FDR < 0.05 versus global FDR >= 0.05. Together, the panels show that topology-derived spatial exposures capture multivariate structural information associated with bulk transcriptional state and recurrent RNA-processing, lipidmetabolic, and tissue-structural programs.

We then tested each topology score against nine prespecified transcriptional modules using the model : module score ∼ 0 + muscle site + trait-specific topology score + Type I fiber fraction.

This model accounts for the exact muscle site and the continuous Type I fiber fraction, allowing topology organization to be evaluated beyond fiber abundance alone. Across the 99 trait-module combinations, the strongest nominal association was a negative relationship between the 24 h pH topology score and the oxidative- mitochondrial module (effect = -0.260, nominal P = 0.0442). Additional directional trends involved vascular/capillary and oxidative-mitochondrial modules for the 24 h meat-color a* score and the oxidativemitochondrial module for 24 h conductivity (Figure 4B; Figure S8). However, no association survived trait-wise or global false-discovery-rate correction. The prespecified modules were relatively small, with 4-7 successfully mapped genes per module. This limited representation may reduce sensitivity to distributed transcriptional changes. We therefore treated the module analysis as an interpretable summary and used genome-wide ranked enrichment as the primary functional follow-up.

For each trait-specific topology score, we ranked retained genes by the moderated t statistic from the primary exact-site model and performed Gene Ontology Biological Process gene-set enrichment analysis using 10,000 permutations and deterministic trait-specific random seeds. Across the 11 traits, 32,101 valid enrichment results were obtained; 412 reached trait-wise FDR < 0.05 and 353 remained significant after global FDR correction across traits and gene sets (Figure 4C).

RNA processing and RNA splicing were the most recurrent functional themes, whereas lipid-metabolic and extracellular-matrix/structural programs appeared in trait-dependent patterns. Because RNA processing, mRNA processing, and RNA splicing occupy overlapping GO levels, we interpret them as one higher-order RNA-processing theme. The recurrence of these programs indicates that multivariate spatial topology is associated with coordinated transcriptional states extending beyond the small marker modules. At the same time, the trait- dependent enrichment directions suggest that different meat-quality phenotypes capture distinct combinations of spatial organization and molecular state.

We evaluated whether module-level associations were sensitive to alternative grouping and sampling definitions by comparing developmental-context adjustment, anatomical-compartment adjustment, and a majormeat-muscle subset with the exact-site model. Effect correlations and sign-agreement rates were generally modest, indicating sensitivity to muscle classification and sample range (Figure S9).

As a further exploratory decomposition, each of the seven raw spatial metrics was tested directly against each of the nine modules without PLS weights or a trait-specific composite score. None of the 63 rawmetric/module combinations reached nominal P < 0.05 and FDR < 0.05 (Figure S10). Thus, the present data do not support assigning the transcriptional associations to a single spatial metric. Instead, the results are most consistent with a multivariate relationship between tissue topology and transcriptional state.

Collectively, these analyses extend the spatial-transcriptomic observations above by showing that topology- derived exposures retain information about bulk transcriptional programs after accounting for muscle site and Type I fiber abundance. The strongest evidence is pathway-level and genome-wide, particularly for RNA processing, lipid metabolism, and tissue-structural programs. The module and raw-metric analyses remain hypothesis-generating, and the current study does not establish a causal effect of topology on transcription or demonstrate that topology is independent of all cellular-composition effects. In the present study, most genes and predefined transcriptional modules did not pass multiple-testing correction, which may be attributable to the tissue-mixture characteristics of bulk RNA-seq data and limited statistical power. Bulk transcriptomes represent the average expression profile of the entire muscle tissue and consequently integrate transcriptional signals from type I and type II muscle fibers, as well as vascular, stromal, immune, and adipose-related cells. In contrast, some of the spatial phenotypes examined in this study primarily reflect type I muscle fiber-related structures. Therefore, type I fiber-specific expression changes may be diluted by signals from other cell types and muscle fiber types. Although the overall proportion of type I fibers was included as a covariate in the models, this adjustment cannot fully disentangle cell type-specific transcriptional effects. In addition, the analysis involved 10,195 genes and multiple phenotypic traits in a dataset comprising 98 samples from 39 muscle sites. The resulting highdimensional testing burden and multiple-testing correction further reduced the statistical power to detect modest effects. Thus, the bulk analysis primarily provides tissue-level directional signals and has limited ability to resolve cell type-specific or spatially localized effects. To address this limitation, we subsequently performed spatial transcriptomic analyses to distinguish the local transcriptional characteristics of different muscle fiber types and other cell populations, and to further evaluate spatial–transcriptional associations that were suggested by, but did not reach statistical significance in, the bulk analysis.

### 5. Myofiber topology links type I fiber organization to local slow-oxidative molecular niches

To focus on type I muscle fibers, we registered myofiber-typing images with spatial transcriptomic data and analyzed spatial gene-expression patterns at two complementary levels to test whether local topology corresponds to slow-oxidative molecular states. First, Global Moran’s I showed significant positive spatial autocorrelation for type I marker genes including MYH7, TNNT1, TNNC1, and TPM3, as well as for slow-oxidative and stress/cytoskeletal modules (Figure 5B). Thus, slow-oxidative expression is not randomly distributed in tissue, but displays global spatial organization. However, global autocorrelation does not imply that all type I fiberenriched regions share the same molecular state. Gene expression can still vary across structurally similar neighborhoods because of local cellular composition, metabolic state, transcriptional activity, and spatial transcriptomic resolution. Therefore, we further used Local Moran’s I to identify local cores of concordantly high expression.

**Figure 5.**
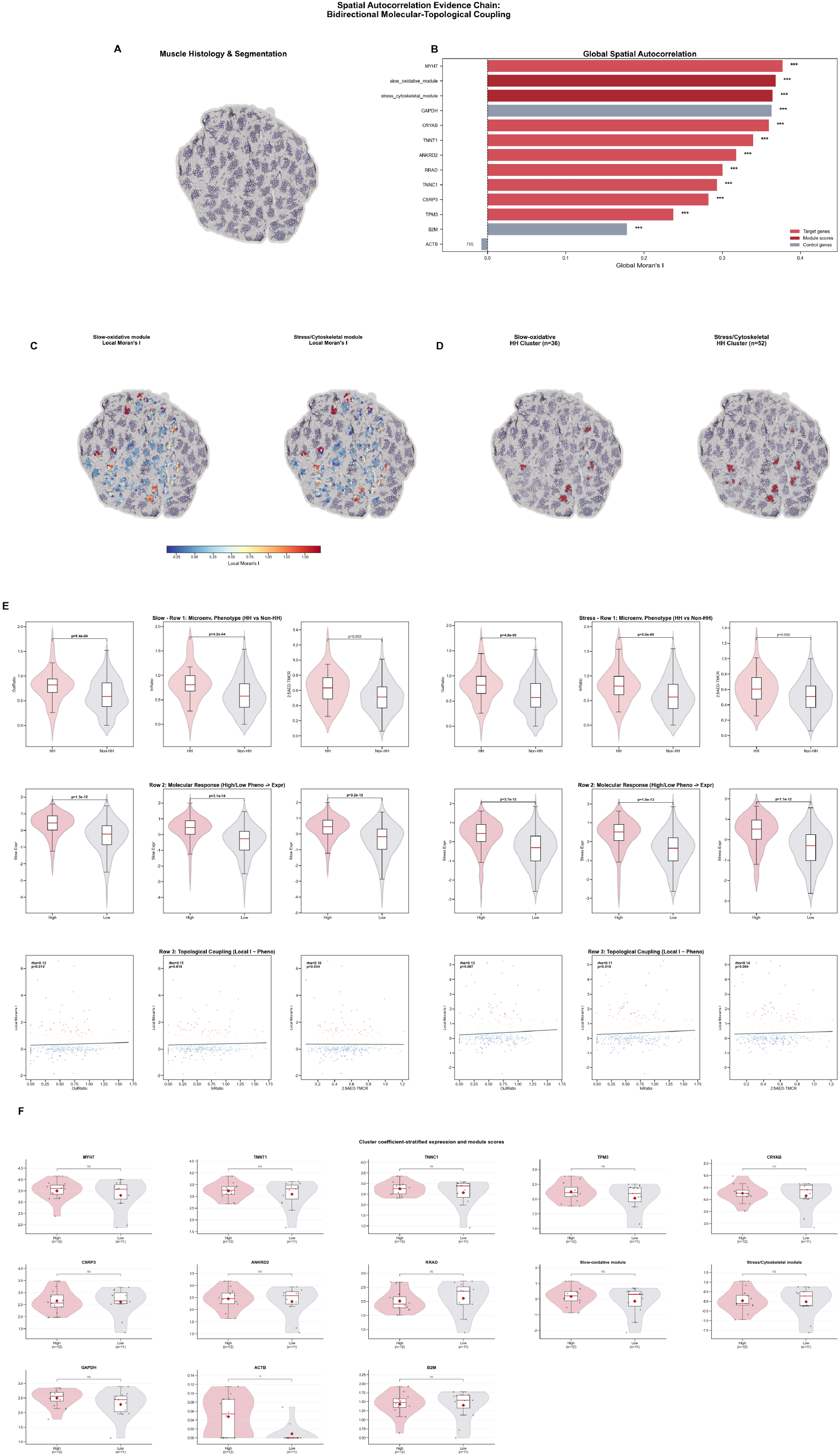
Spatial autocorrelation links type I myofiber topology with slow-oxidative molecular programs.(A) PMM muscle section with segmented myofiber contours. (B) Global Moran’s I analysis showing significant spatial autocorrelation of type I fiber marker genes and slow-oxidative/stress-cytoskeletal module scores. (C) Local Moran’s I maps of the slow-oxidative and stress/cytoskeletal modules. (D) High-high hotspot regions identified for the two modules. (E) Statistical validation of the associations among hotspot status, type I fiber neighborhood metrics, module expression, and local spatial autocorrelation (including OutRatio, InRatio, and 2.5AEDTMCR). (F) Stratified comparison (by cluster coefficient) of target gene expression, module scores, and control gene expression.

Local Moran’s I analysis showed that slow-oxidative and stress-cytoskeletal modules formed high-high (HH) hotspots only in subsets of the tissue, where both the target position and its spatial neighbors showed high expression with spatial statistical significance (Figure 5C,D). Mapping these HH hotspots back to the registered myofiber images showed that: InRatio, OutRatio, and 2.5AED-TMCR were significantly higher in HH regions than in non-HH regions; consistently, the clusters with tighter intra-connections also exhibit higher slow- oxidative/stress-cytoskeletal module scores (Figure 5E,F). High type I fiber adjacency and local enrichment therefore provide a structural background for spatial aggregation of slow-oxidative molecular programs. In other words, myofiber topology defines potential slow-oxidative neighborhoods, whereas Local Moran’s I further identifies slow-oxidative niches with concordantly enhanced expression. Gene-level correlations supported this interpretation. In PMM, MYH7, TNNT1, and TNNC1 were positively correlated with 2.5AED-TMCR, InRatio, and OutRatio (Figure S11C). Genes correlated with these topology metrics also included CRYAB, CSRP3, ANKRD2, and RRAD, which are related to oxidative metabolism, mechanical adaptation, and metabolic homeostasis. Similar trends were observed in SM: TNNC1 was positively correlated with betweenness centrality, whereas the type II fiber marker MYL1 was negatively correlated with InRatio (Figure S11D).

These findings are consistent with recent studies of porcine skeletal muscle spatial microenvironments. MYH7-marked spatial domains have been reported to form aerobic-metabolism-associated microenvironments involving myofibers, fibro/adipogenic progenitors, immune cells, and stromal cells [86]. Our results further show, from the perspective of myofiber topology, that local contact, clustering, and enrichment of type I fibers can serve as structural cues for identifying slow-oxidative microenvironments. Together with reports that clustered type I fibers enlarge in aged human muscle [30], these results suggest that myofiber topology may represent a conserved phenotypic dimension linking tissue architecture, metabolic state, and muscle adaptation.

Collectively, type I myofiber spatial topology is not merely a passive consequence of fiber abundance. Rather, it is a tissue-structural feature coupled to single-fiber phenotype and local slow-oxidative molecular programs. These image-based and spatial-transcriptomic results establish the molecular meaning of myofiber topology and motivate the following analyses of the separability of type I fiber abundance and topology organization capacity, as well as topology remodeling across muscle sites and breeds.

### 6. Type I myofiber abundance and spatial topology organization capacity are separable structural dimensions

The preceding results indicate that local adjacency and clustering of type I fibers have molecular meaning. However, many spatial metrics naturally change with type I fiber abundance: when a tissue contains more type I fibers, they are more likely to approach each other and form local enrichment. Direct comparison of raw spatial metrics may therefore confound increased fiber number with enhanced spatial topology. To determine whether spatial topology constitutes a phenotype dimension independent of abundance, we separated type I fiber abundance from topology organization capacity within each breed.

Representative comparisons suggested that type I fiber proportion and mean area alone do not fully describe tissue structure. In LW pigs, TBLO and TIC showed limited differences in type I fiber proportion and area, but differed markedly in cluster number, inter-cluster distance, clustering coefficient, and global distance entropy (FigureS2). Similarly, muscles with similar type I fiber proportions could display either multiple dispersed small clusters or more compact and continuous local connected structures. Thus, similar fiber abundance can be spatially implemented as distinct microenvironmental architectures.

To quantify this phenomenon systematically, we built within-breed abundance-adjustment models using Ratio of Stained Cells to predict each topology metric. The observed deviation from the model expectation was then direction-corrected (Table S5) and standardized. We integrated multiple abundance-adjusted topology metrics into a Topology Enhancement Score (TES) (Figure 6A-D). TES > 0 indicates that a muscle exhibits stronger-than-expected spatial organization at a given type I fiber proportion, whereas TES < 0 indicates weakerthan-expected organization.

**Figure 6.**
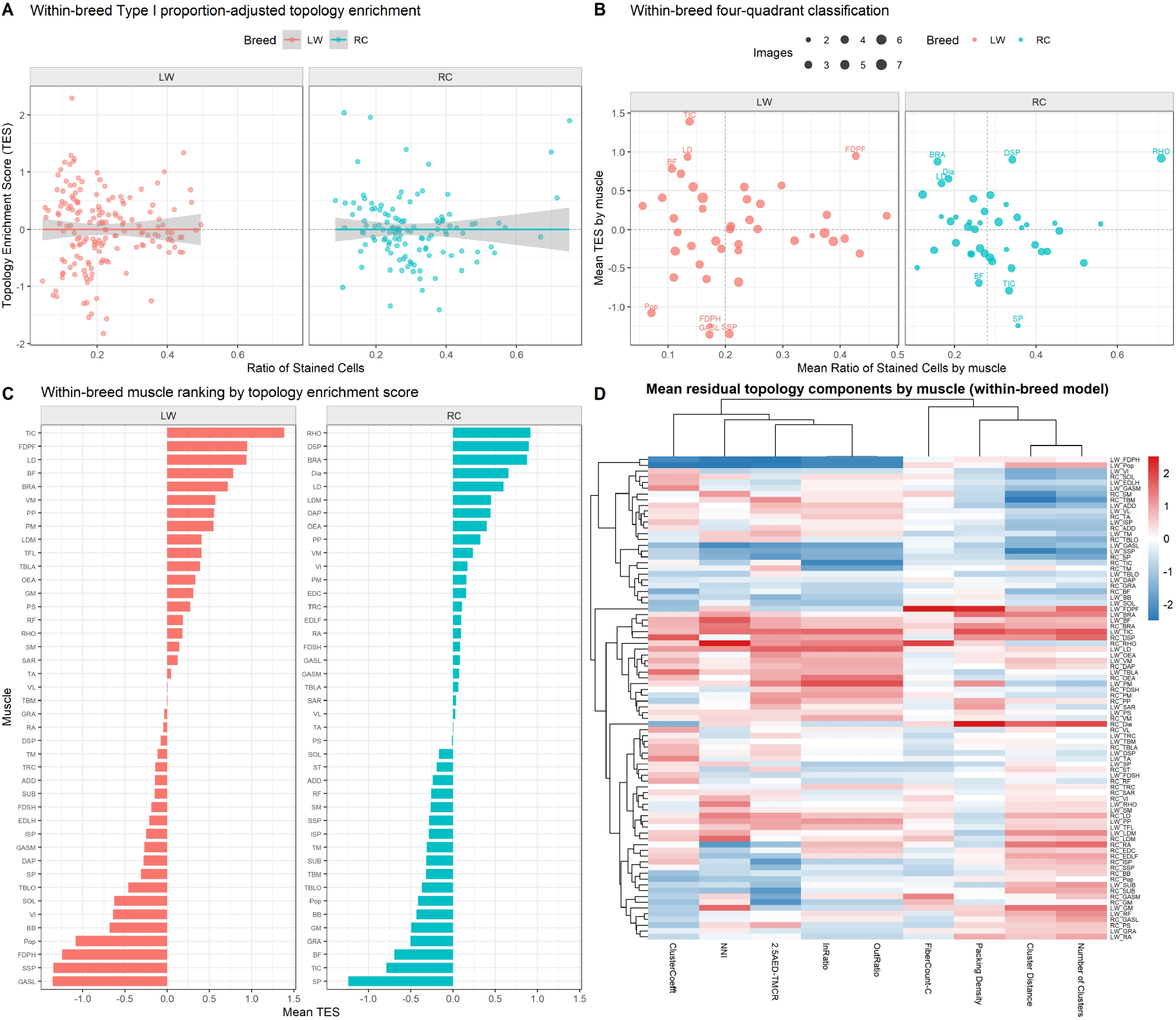
Abundance-adjusted Topology Enhancement Score distinguishes type I fiber abundance from spatial topology organization capacity. (A) The relationship between the proportion of type I muscle fibers and the topological reinforcement score (TES) in LW and RC pigs. Each point represents an image sample. The solid line and shading represent the linear fitting trend and its confidence interval. (B) Quadrant classification based on the average proportion of type I muscle fibers and the average TES of the muscles. The x-axis represents the average proportion of type I muscle fibers of the muscle, and the y-axis represents the average TES of the muscle; the size of the points indicates the number of image samples included in the analysis for each muscle, and the dotted lines represent the reference boundaries on the corresponding axes. (C) The results of the ranking of each skeletal muscle by average TES in LW and RC pigs. Positive values indicate that the muscle has a relatively higher proportionally corrected topological reinforcement level within the breed, while negative values indicate a relatively lower one. (D) The residual topological composition heatmap of each muscle obtained within the breed. The list represents the spatial topological indicators used to construct TES, the rows represent the breed-muscle combinations; the color indicates the standardized residual level, red represents higher than the model expectation, and blue represents lower than the model expectation.

TES analysis showed that type I fiber abundance and spatial topology capacity are not equivalent (Figure 6A,B). Within LW and RC pigs, some muscles showed high TES despite relatively low type I fiber proportions, indicating that they can form strong local organization from a limited type I fiber basis. Conversely, some muscles had high type I fiber proportions but low TES, suggesting that increased fiber number does not necessarily imply enhanced topology. For example, in LW, TIC represented a low-abundance/high-topology muscle, whereas FDPF displayed both high type I fiber abundance and high TES, representing a high-abundance/high-topology strategy (Figure 6B). When viewed at the overall body level, there may be systematic differences in the spatial organization strategies of muscle fibers among different breeds (Figure 6C).

The components of TES further showed that abundance-adjusted topological identity is not driven by a single metric (Figure 6D). In others, it reflected combined contributions from within-cluster fiber count, nearestneighbor index, packing density, inter-cluster distance, and 2.5AED-TMCR. Different skeletal muscles can therefore reach similar TES levels through different topological routes, indicating that spatial organization capacity is intrinsically multidimensional. In some muscles, high TES was mainly contributed by stronger within- cluster connectivity or higher clustering coefficients.

This result moves the description of skeletal muscle from “what fiber types are present” to “how fibers are organized.” Type I fiber proportion indicates how many slow-oxidative-related fibers are present in the tissue, whereas TES describes whether these fibers form tighter, more enriched, or more complex spatial organization at a given abundance. Together, they define the slow-oxidative structural state of skeletal muscle: similar type I fiber proportions can correspond to different local adjacency, cluster expansion, and global distribution patterns, and similar topology levels can be achieved through different combinations of spatial metrics.

In summary, type I myofiber spatial topology is not a simple extension of conventional proportion and area metrics, but a tissue-structural dimension separable from fiber abundance. Establishing this dimension enables two further questions: whether different skeletal muscles have stable and distinguishable topological identities, and whether breed background remodels these identities in a muscle-specific manner.

### 7. Distinct skeletal muscles have distinguishable type I fiber spatial topology identities

Having shown that type I fiber proportion and topology organization capacity are separable, we next reexamined the high-dimensional phenotype space of the LW multi-muscle dataset to determine whether different skeletal muscles carry stable spatial topology identities. Here, “topology identity” refers to a tissue-structural state jointly defined by type I fiber abundance, morphology, local adjacency, connected-cluster architecture, and global distribution, rather than by any single proportion or area metric.

Unsupervised hierarchical clustering showed that skeletal muscle samples did not simply group according to anatomical regions such as trunk, forelimb, or hindlimb. Instead, they formed phenotypic subtypes that crossed anatomical boundaries (Figure 7A D). In phenotypic space, some trunk muscles were close to each other, whereas other trunk muscles clustered with limb muscles. This cross-regional convergence suggests that type I fiber spatial organization is more likely related with the contractile tasks, metabolic demands, and local tissue environment of each muscle than anatomical position itself.

**Figure 7.**
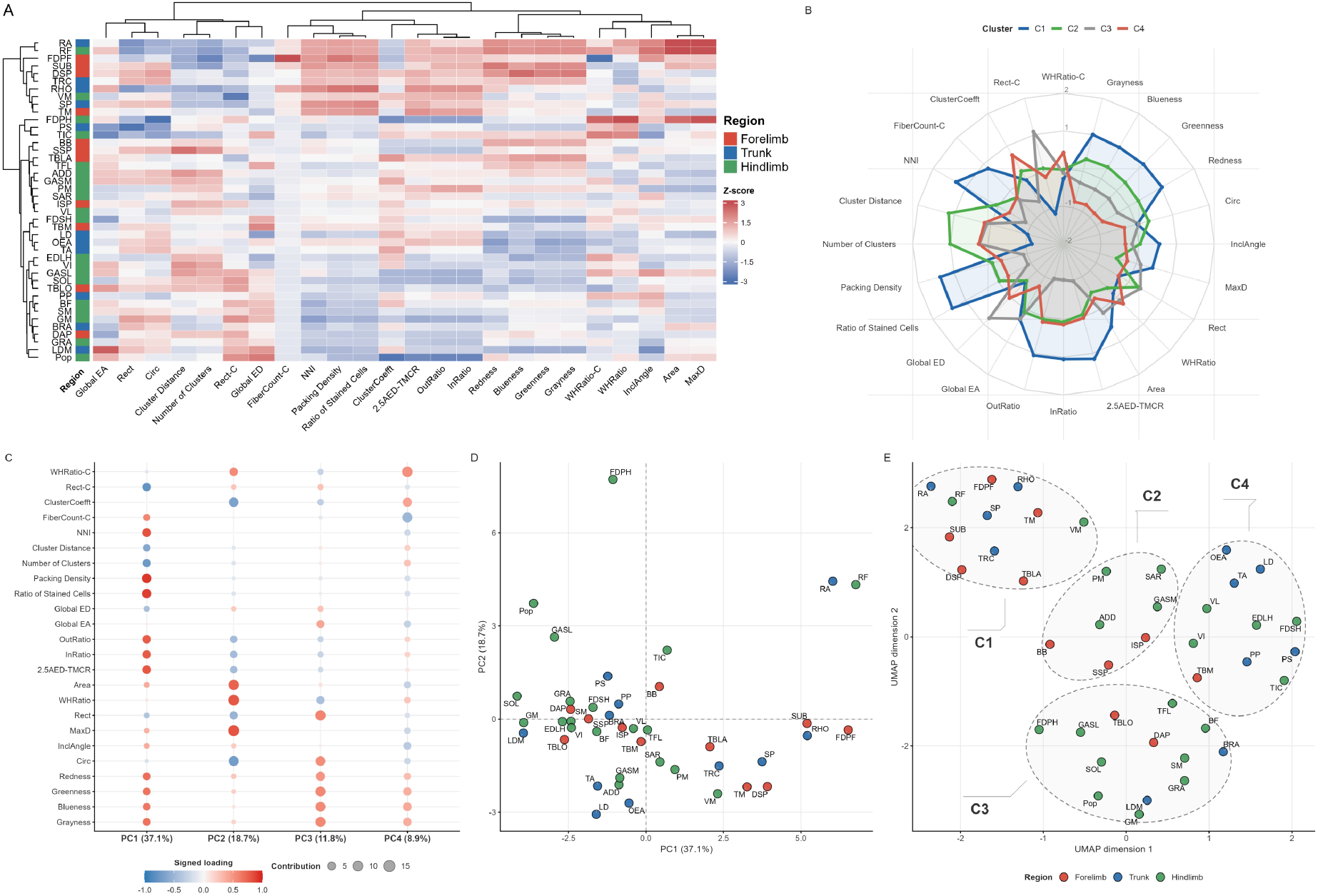
High-dimensional structure of spatial phenotypes across LW skeletal muscles. Hierarchical clustering, PCA, and UMAP show that skeletal muscles do not simply group by anatomical region but form distinguishable spatial phenotypic subtypes. (A) Unsupervised hierarchical clustering heatmap based on all phenotypic indicators. (B) The mean distribution of the four spatial phenotypic subtypes (C1-C4) on each phenotypic index. (C) Principal component analysis (PCA) scatter plots of all phenotypic indicators and the load distribution of each principal component. (D) Score distribution plots of each skeletal muscle in the PC1 and PC2 dimensions. (E) Two-dimensional embedding graph of UMAP based on principal components of PCA.

PCA decomposed multi-site differences into interpretable dimensions. PC1 was mainly contributed by global distribution, adjacency, and local enrichment metrics; PC2 reflected fiber geometry; PC3 corresponded to SDH staining intensity; and PC4 was related to connected-cluster and aggregation structure (Figure 7C). Because both PC1 and PC4 were dominated by spatial organization features, topology is not merely accessory information to fiber proportion or area, but one of the major phenotypic axes distinguishing skeletal muscles.

Using UMAP embedding and K-means clustering, LW skeletal muscles could be summarized into four spatial phenotypic subtypes (Figure 7E): C1, a high-density/strong-connection subtype with higher type I fiber proportion, local adjacency, and staining intensity; C2, a dispersed small-cluster subtype with many small connected clusters; C3, a low-density/high-heterogeneity subtype with high global entropy and weak connectivity; and C4, a morphology-driven subtype defined mainly by larger or elongated fibers (Figure 7B). Thus, skeletal muscles are not arranged along a single axis of “more or fewer slow fibers”, but form distinct tissue states through multiple spatial organization strategies.

Representative comparisons illustrate the biological meaning of these topology identities. LD, TA, and RA are all trunk-related muscles, but LD and TA were more morphology-driven, whereas RA showed higher type I fiber density, stronger adjacency, and stronger SDH staining (Figure S12A). TBLA and BB are both forelimb muscles and have similar conventional type I proportion, area, and morphology metrics, yet TBLA showed higher clustering coefficient and adjacency, indicating a stronger tendency to form compact local slow-oxidative units (Figure S12B). Similarly, RHO and FDPF had comparable type I fiber proportions, but RHO contained larger connected clusters and stronger adjacency (Figure S12C). These examples show that similar abundance can be organized into distinct local microenvironments.

At a finer site-comparison level, spatial metrics also captured structural specialization that conventional metrics had difficulty separating. TBLA, TBM, and BB differed only modestly in type I fiber proportion and area, but showed gradients in adjacency, cluster shape, and clustering coefficient (Figure S12D). LDM, PS, and SM also displayed distinct global distributions and local connection patterns despite similar type I fiber proportions (Figure S12E). Even among anatomically adjacent triceps brachii subregions, spatial topology distinguished microstructural specialization (Figure S12F). These skeletal muscles have been shown to differ in structure, metabolism, and quality-related traits [87-90], but the key point here is that site differences involve not only fiber-type composition or size, but also how type I fibers are arranged within tissue. Previous studies that only explained macroscopic phenotypic differences (metabolism, quality-related traits and bulk transcriptome characteristics) based on muscle fiber proportions and areas, may have led to inaccurate conclusions.

Together, different skeletal muscles have distinguishable type I fiber spatial topology identities. These identities are jointly defined by abundance, morphology, local adjacency, connected clusters, and global distribution, with spatial organization features representing major sources of variation. This site-level topology atlas provides a basis for testing whether breed background remodels these identities and whether such remodeling is related to tissue-level traits.

### 8. Breed background reveals both conservation and plasticity of type I fiber spatial topology identity

Section 6 showed that different muscles have distinguishable type I fiber spatial topology identities. To test whether these identities are stable and whether they are modulated by genetic or breed background, we compared multiscale type I fiber phenotypes between LW and RC pigs across corresponding skeletal muscle sites. Breed background was treated as a systematic perturbation that reveals which structural components of muscle-site identity are conserved and which are plastic.

Both LW and RC pigs showed clear type I fiber clustering. In both breeds, the average number of fibers per connected cluster exceeded seven, and more than 90% of type I fibers were located within clusters containing at least two fibers. Thus, type I fiber clustering is a basic organizational feature shared across breeds. At the same time, the breeds showed systematic differences: RC had higher Ratio of Stained Cells, Packing Density, 2.5AEDTMCR, and ClusterCoeff, whereas LW had larger and more regular type I fibers (Figure 8A). Notably, breed differences in InRatio and OutRatio were not prominent at the global mean level, suggesting that direct adjacency strength does not shift uniformly across all muscles and may be regulated by site-specific factors.

**Figure 8.**
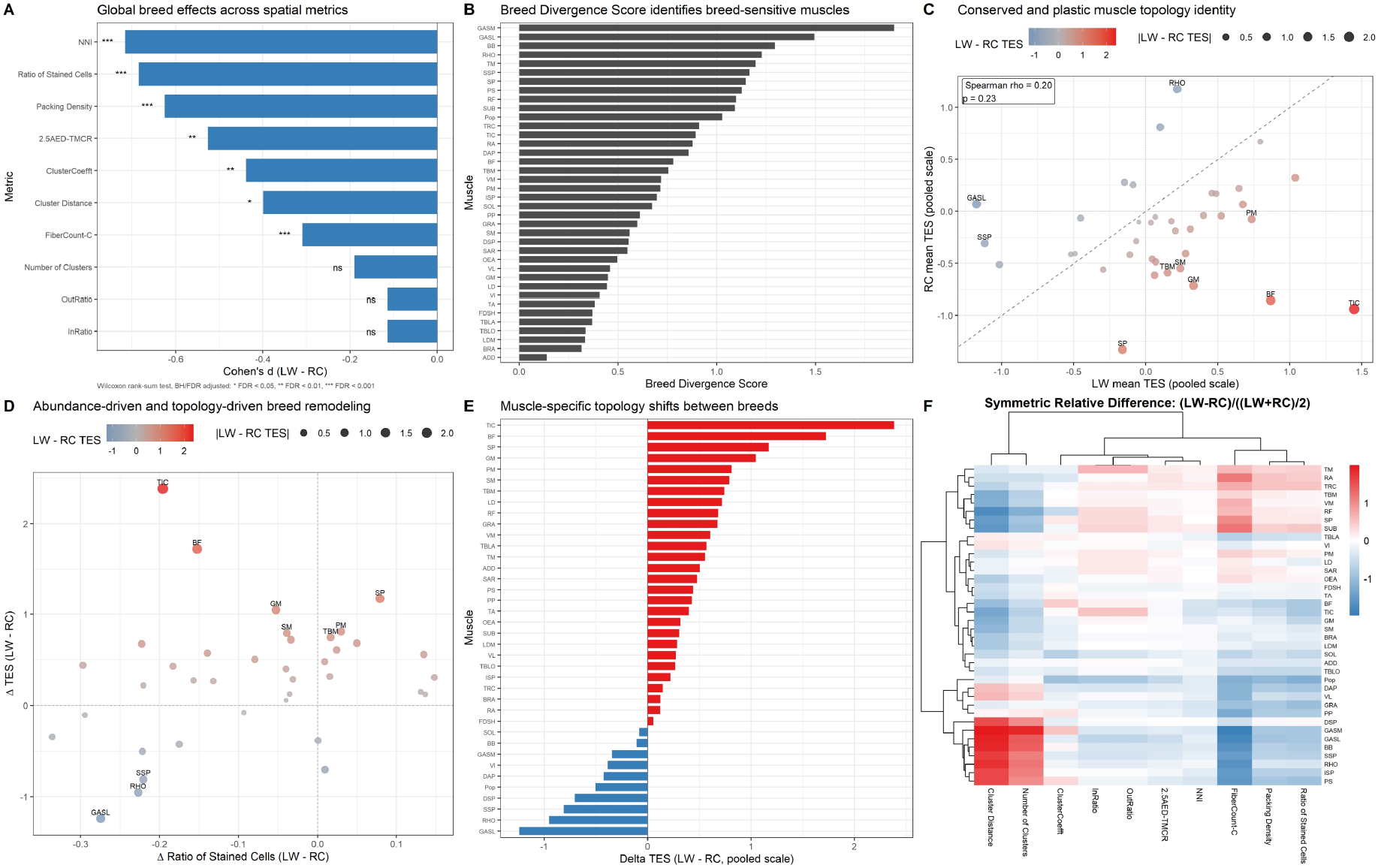
Effects of breed background on global spatial metrics and TES. Breed Divergence Score identifies skeletal muscles that are more sensitive to breed background. (A) The magnitude of breed effect for global spatial phenotypic indicators between LW and RC pigs. The bar chart represents Cohen’s d (LW - RC), and the significance is based on the Wilcoxon rank sum test and corrected by BH/FDR. (B) Breed Divergence Score calculated based on multi-index differences, used to identify skeletal muscles more sensitive to the breed background. The higher the score, the stronger the spatial phenotypic differentiation of the muscle between LW and RC. (C) Comparison of the average TES of the shared muscles between LW and RC. The dotted line represents the reference line where the TES of the two breeds are completely consistent; the point color indicates the TES difference between LW and RC, and the point size indicates the absolute value of the TES difference between the two breeds. Spearman correlation is used to evaluate the conservation of muscle topological identity between the two breeds. (D) Joint analysis of the difference in I-type muscle fiber proportion and TES difference. The x-axis is Δ Ratio of Stained Cells (LW - RC), and the y-axis is ΔTES (LW - RC); the color and size of the points indicate the direction and magnitude of the TES difference, used to distinguish abundance-driven and topological remodeling type breed differences. (E) Ranking of ΔTES (LW - RC) for each skeletal muscle. Red indicates that the muscle has a higher TES in LW, and blue indicates that it has a higher TES in RC, used to show the direction of topological reinforcement between the breeds. (F) Symmetric relative difference heatmap of spatial phenotypic indicators between LW and RC, calculated as (LW - RC) / ((LW + RC) / 2). Red indicates that LW is higher than RC, and blue indicates that RC is higher than LW, used to show the muscle-specific remodeling patterns of specific topological indicators.

To compare topological identities on a common scale, we used a pooled model to adjust for type I fiber proportion and recalculated TES for homologous muscles. If a muscle shows similar TES in both breeds, its topology identity is relatively conserved; if it deviates substantially, its type I fiber spatial organization has been remodeled by breed background (Figure 8B). The TES correlation between shared LW and RC muscles was weak and not statistically significant (Spearman ρ = 0.20, *p* = 0.23; Figure 8C), indicating that muscle topology identity is not highly conserved between the two breeds.

This inconsistency was not random noise but showed clear muscle specificity. A Breed Divergence Score based on multimetric differences identified GASM, GASL, BB, RHO, TM, SSP, SP, PS, and RF as muscles with relatively high breed divergence, whereas other muscles maintained more stable spatial structures (Figure 8B,C). Therefore, breed background does not impose a single global effect on whole-body skeletal muscle. Instead, it selectively remodels type I fiber spatial organization according to muscle-site identity and tissue context.

Joint analysis of *Δ* Ratio of Stained Cells and *Δ* TES further separated the sources of breed differences (Figure 8D,E). Some muscles showed concordant changes in type I fiber abundance and TES, suggesting abundance increase accompanied by topology enhancement. Other muscles showed clear *Δ* TES despite similar type I fiber proportions, indicating that breed background mainly altered the spatial organization mode rather than simply changing the number of type I fibers. For example, VI and DSP had similar type I fiber proportions between breeds, but LW showed more fiber clusters and greater inter-cluster distances, suggesting that the same abundance can be implemented as more dispersed or more continuous spatial arrangements (Figure S13, Figure S14).

Metric-level comparison further supported a breed-by-site interaction. LW showed higher type I fiber proportions only in SP, TRC, RA, RF, SUB, TM, and a few other sites, whereas most other muscles had higher proportions in RC. Except for DSP, ISP, LD, and several other sites, LW generally had larger type I fibers (Figure S13, Figure S14A,B). However, proportion and area could not explain all differences. The ranking of metrics across muscles often differed between LW and RC, especially for within-cluster fiber number, local adjacency, and global distribution metrics (Figure 9A-E). More specifically, traditional metrics such as proportion and crosssectional area are the most conservative, while spatial distribution and SDH staining intensity show more remarkable systematic differences. PERMANOVA showed that breed explained 12.8% of total phenotypic variation (*R*^*2*^ = 0.128, *p* = 0.001) and 11.2% of variation in global spatial-distribution metrics. PCA and UMAP also showed a separation tendency between breeds (Figure 9F,G). These results indicate that breed background leaves a stable spatial phenotypic signal at the cross-site scale, but the signal is expressed through muscle-specific remodeling.

**Figure 9.**
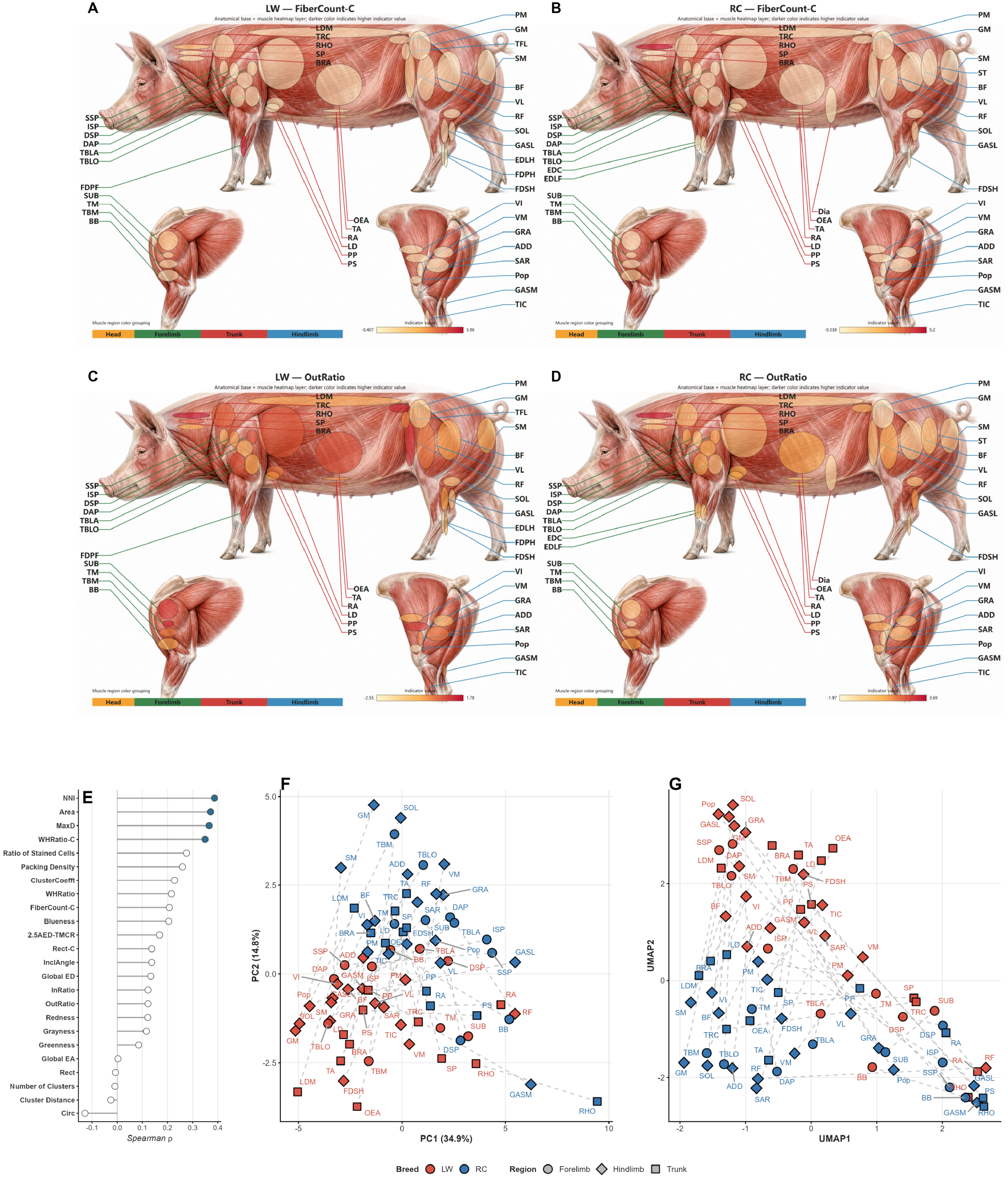
Within-breed and between-breed spatial phenotype structure. Different skeletal muscles show inconsistent rankings across within-cluster fiber number, adjacency, and global distribution metrics. (A-D) Special indicators show differences in the internal expression of pigs, and the skeletal muscle tissue structure is inconsistent. (E) Phenotypic correlations among pig breeds. (F-G) The PCA scatter plot distribution of the samples and the UMAP two-dimensional embedding distribution.

This muscle-specific remodeling provides an entry point for understanding tissue-level traits. Previous studies reported that differences in pH, lightness, and marbling between lean-type pigs and indigenous pigs are more pronounced in LDM than in SM [91]. In this study, LW and RC did not differ significantly in type I fiber proportion in LDM, but RC had stronger clustering and adjacency features. In contrast, spatial phenotype differences in SM were smaller (Figure S13G, Figure S14). Site-specific topological remodeling may therefore be closer to the structural background of tissue-trait variation than whole-body average differences in type I fiber proportion.

In summary, differences between LW and RC are not limited to type I fiber proportion or single-fiber size. They reflect both conservation and plasticity of skeletal muscle topology identity. Type I fiber clustering is a shared basic structure across breeds; on this basis, different muscles show varying degrees of abundance change, topology remodeling, or both. This result moves breed comparison from single-metric analysis to system-level spatial phenotype remodeling.

### 9. Automated instance segmentation supports scalable myofiber spatial phenotyping

Because all topology metrics depend on accurate instance-level fiber contours, scalable analysis of SDH-positive type I myofibers required a robust segmentation workflow. We therefore developed an improved Mask R-CNN framework [65], trained with SAM-assisted and Labelme-corrected annotations, to handle the dense, small, irregular, variably stained, and frequently adjacent morphology of swine muscle fibers. The model (Figure 10A and B) incorporated modulated deformable convolution (DCNV2) for adaptive feature extraction [48], Soft-NMS for crowded-instance post-processing [49], normalized weighted distance (NWD) loss for smallobject localization [92], test-time augmentation (TTA) for orientation and scale robustness [87], and automatic mixed precision (AMP) training to improve computational efficiency [88].

**Figure 10.**
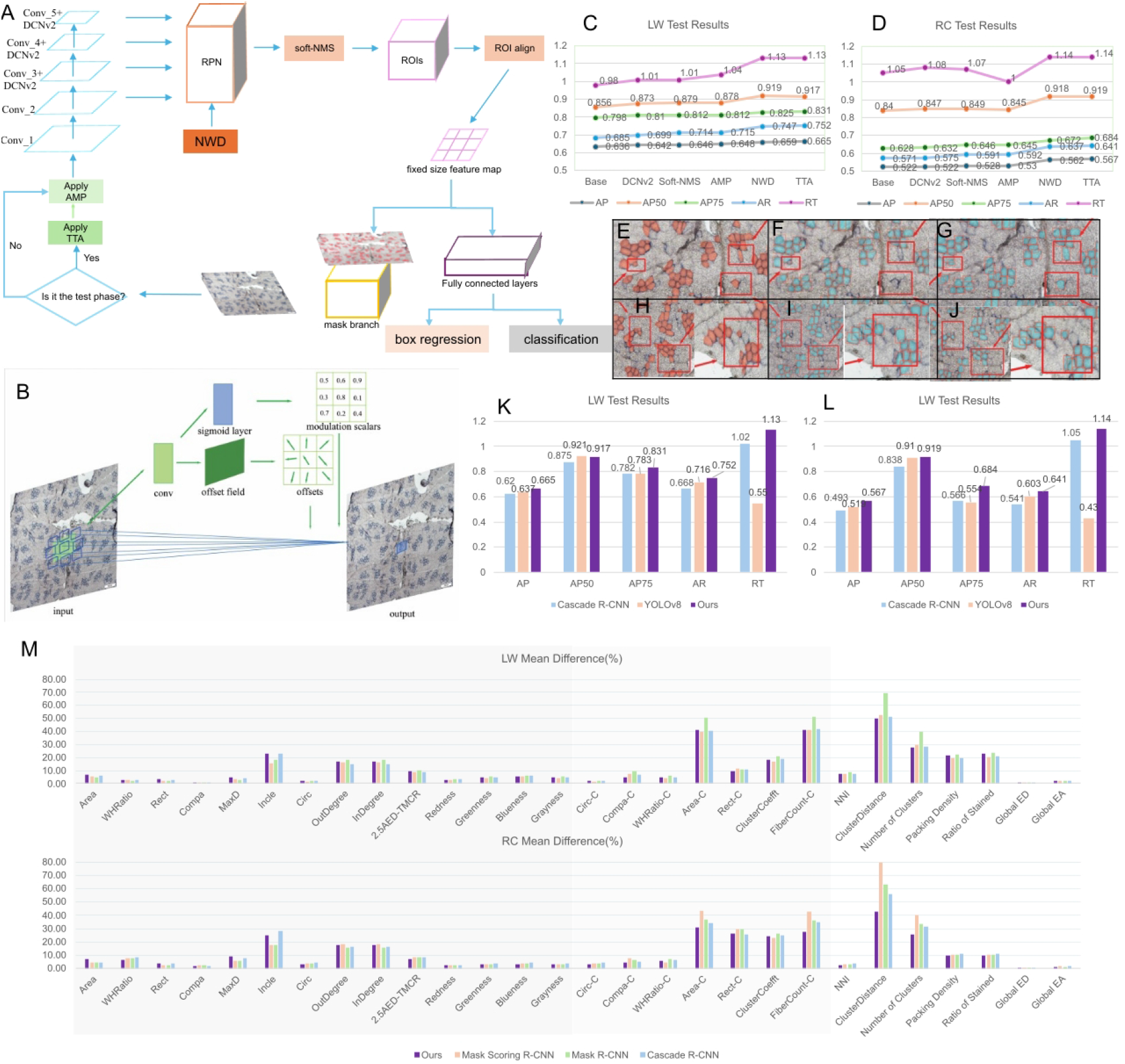
Improved Mask R-CNN supports scalable myofiber instance segmentation and downstream spatial phenotype extraction. The overall framework of the improved Mask R-CNN. (B) The modulated deformable convolution process. (C-D) Ablation study results of Mask R-CNN, where “Base” means the original Mask R-CNN. (E) Manually annotated image. (F) The predicted image after corresponding to NMS. (G) The predicted image corresponding to soft-NMS. (H) The manually annotated image. (I) The predicted image corresponding to standard IoU. (J) The predicted image corresponding to NWD. (K-L) Comparison experiment results with other models. (M) The differences between phenotypic values from manually annotated myofiber contours and those derived from model-automated segmentation contours.

Ablation analyses showed that these additions improved segmentation accuracy in both LW and RC datasets. Relative to baseline Mask R-CNN, the improved model increased AP_50_ by 6.1% and AR by 6.7% in LW, and increased AP_50_ by 7.9% and AR by 7.0% in RC (Figure 10C-D). Soft-NMS, NWD, and related techniques effectively reduced missed detections of muscle fibers (Figure 10E-J). Benchmark comparisons with Cascade RCNN [83] and YOLOv8[93] further showed overall advantages in AP, AP_50_, AP_75_, AR and RT (Figure 10K,L), indicating that the optimized workflow better preserved irregular contours and crowded neighboring fibers.

We next asked whether this performance was sufficient for biological phenotyping. Predicted contours and manual annotations were processed through the same multiscale feature-extraction pipeline. Single-fiber morphology and SDH-intensity features generally showed mean errors below 10%, and global distribution metrics such as NNI and global compactness were also mostly within 10% (Figure 10M). Topology-derived features, including connected-cluster size and inter-cluster distance, were more sensitive, with errors of approximately 20%-70%. Thus, the automated workflow supports scalable extraction of major morphometric and global spatial signals, while topology-sensitive readouts should continue to be evaluated by phenotype-level error propagation rather than AP or IoU alone.

## DISCUSSION

### Spatial topology as a phenotype layer beyond fiber abundance

This study establishes type I myofiber spatial topology as a quantitative phenotype layer of skeletal muscle. Traditionally, muscle metabolic properties have been interpreted mainly through fiber-type composition, fiber size, and histochemical intensity. Our results show that these descriptors capture only part of the tissue phenotype. By measuring single-fiber morphology and staining, local contact, connected-cluster architecture, diffusion-ring neighborhoods, and global tissue configuration, we found that muscles with comparable type I fiber proportions can exhibit markedly different spatial organizations. Thus, type I myofiber abundance and spatial organization capacity are related but non-equivalent properties.

A key technical requirement for this framework is reliable instance-level segmentation. SDH-positive type I fibers are often small, densely packed, irregularly shaped, variably stained, and frequently adjacent, making manual annotation and conventional segmentation difficult to scale. We therefore developed an improved Mask R-CNN workflow based on SAM-assisted and Labelme-corrected annotations, with modulated deformable convolution, Soft-NMS, normalized weighted distance, test-time augmentation, and automatic mixed precision training. These modifications improved detection and segmentation performance, particularly by reducing missed fibers in crowded regions. Importantly, this also highlights a broader methodological point: topology phenotypes depend not only on pixel-level accuracy, but also on whether segmentation errors alter neighborhood, clustering, and connectivity readouts. Future image-analysis pipelines for tissue topology should therefore evaluate phenotype-level stability in addition to conventional segmentation metrics.

### Neighbor-dependent clustering and local slow-oxidative niches

Our data support a neighbor-dependent model of type I myofiber organization. The topology enhancement score separated muscles that merely contained many type I fibers from those that formed stronger-than- expected local connectivity at a given abundance. This distinction is biologically meaningful because local clustering can create microenvironments with different metabolic, mechanical, and signaling properties from those inferred by fiber proportion alone. In this sense, type I fibers should be viewed not only as individual metabolic units, but also as components of local tissue neighborhoods.

Spatial transcriptomic integration further connected these structural neighborhoods with molecular organization. Slow-oxidative genes and functional modules, including markers related to contractile identity, oxidative metabolism, cytoskeletal organization, and stress adaptation, showed non-random spatial patterns. High-expression domains tended to overlap with regions enriched for type I fiber contact and neighborhood- based topology metrics. These findings suggest that spatially clustered type I fibers may correspond to molecularly active slow-oxidative niches. However, the relationship should be interpreted as spatial association rather than direct causality, because the current data do not establish whether fiber topology drives local transcriptional states, reflects them, or emerges together with them during muscle development and remodeling.

### Muscle identity and breed-dependent remodeling

Across more than 40 skeletal muscle sites, topology features revealed muscle identities that were not fully explained by anatomical position or type I fiber abundance. Some muscles formed compact and highly connected type I fiber networks, whereas others showed more dispersed or weakly connected patterns. The contribution of spatial features to major phenotypic axes indicates that skeletal muscles are not arranged along a simple “more versus fewer slow fibers” continuum. Instead, each muscle appears to occupy a multidimensional topology state defined by abundance, morphology, neighborhood contact, clustering, and global distribution.

Breed background further remodeled these topology states in a muscle-specific manner. Large White and Rongchang pigs shared some common organizational principles, but the correspondence of topology enhancement across matched muscles was weak, and breed explained a measurable fraction of total phenotypic variation, especially for global spatial-distribution metrics. These results suggest that genetic background, production history, and muscle-specific developmental programs may jointly shape the spatial architecture of slow-oxidative fibers. Because the present study is observational, these breed effects should not be interpreted as direct consequences of selection alone. Controlled developmental, genetic, and intervention studies will be needed to determine how stable or plastic these topology states are.

### Implications for meat quality

The association between topology features and meat-quality traits suggests that spatial organization contributes information beyond conventional fiber composition. In samples with matched phenotypes, topology-aware models improved the explanation of pH, color, conductivity, and intramuscular fat, and removing spatial metrics reduced predictive performance. Features related to connected clusters, local contact, and diffusion-ring neighborhoods were repeatedly selected as important predictors. These findings imply that meat quality may depend not only on how many oxidative fibers are present, but also on how they are arranged within the tissue.

This interpretation is consistent with the idea that local fiber neighborhoods influence postmortem metabolism, water-holding properties, light scattering, and lipid deposition through spatially structured cellular and extracellular environments. Nevertheless, the predictive models should be viewed as phenotype-association models rather than mechanistic proof. Their value lies in showing that topology is not redundant with fiber proportion or area, and that spatial descriptors can provide an additional layer for understanding muscle-tomeat conversion.

### Limitations and future directions

Several limitations should be considered. First, the framework focuses on SDH-positive type I fibers and does not yet jointly model the full spectrum of fiber types, extracellular matrix, vasculature, nerves, and adipocytes. Integrating multiplex histology, immunostaining, and spatial omics would allow a more complete view of muscle tissue organization. Second, although the improved deep learning pipeline substantially increased scalability, topology metrics remain sensitive to segmentation errors in dense or ambiguous regions. Future work should include uncertainty-aware segmentation, cross-laboratory validation, and direct benchmarking of topology stability under annotation and imaging variability. Third, the spatial transcriptomic analysis links tissue architecture to molecular domains, but higher-resolution and perturbation-based studies are needed to resolve causality.

Overall, this study proposes a multiscale framework for modeling skeletal muscle as a spatially organized tissue system. By combining robust deep learning-based segmentation, quantitative topology metrics, spatial transcriptomics, breed comparison, and meat-quality association, we show that type I myofiber topology represents an interpretable and measurable phenotype layer beyond fiber abundance. This perspective may help connect muscle structure, molecular state, genetic background, and product quality in a unified spatial framework.

## RESOURCE AVAILABILITY

### Materials availability

This study did not generate new unique biological reagents. Muscle tissue samples were used mainly for cryosectioning, SDH staining, image acquisition and Spatial transcriptome sequencing. Availability of remaining tissue materials depends on residual sample status and institutional ethical and data-sharing regulations.

### Data and code availability

The datasets generated in this study are available from the corresponding author upon reasonable request. Public deposition of the relevant datasets is planned after completion of the ongoing analyses.

### Author Contributions

Conceptualization, Z. S., Y. J., M. L., and L. J.; methodology, Z. S., Y. J., and G. H.; software, Y. J. and C. G.; formal analysis, Z. S. and Y. J.; data curation, G. Z. and J. C.; writing-original draft, Z. S., and Y. J.; writing-review and editing, M. L. and L. J.; visualization, Z. S., and Y. J.; supervision, Z. S., M. L., and L. J.; project administration, Z. S., M. L., and L. J.; resources, Y. J., and M. L.

## DECLARATION OF INTERESTS

The authors declare no competing interests.

## STAR★METHODS

**KEY RESOURCES TABLE**

See Table S2.

## EXPERIMENTAL MODEL AND SUBJECT DETAILS

### Animals, breeds, and skeletal muscle sampling

This study included two skeletal muscle sample batches. The first batch was collected from four female Rongchang pigs (RC; approximately 10 months old, ∼75 kg) and four female Large White pigs (LW; approximately 10 months old, ∼153 kg). It covered 46 skeletal muscle sites representing major forelimb, trunk, and hindlimb muscles. Full muscle names, abbreviations, and anatomical regions are provided in Table S3. This batch was used to construct the multiscale spatial phenotype system for type I myofibers and to support cross- muscle, cross-breed, neighbor-dependent clustering, TES, and spatial transcriptomic analyses.

The second batch was collected from 17 Landrace pigs (LR) and 20 Large White pigs (LW), including psoas major muscle (PMM) and longissimus dorsi muscle (LDM). These samples were subjected to both meat-quality measurements and corresponding SDH-stained cryosection preparation. Each meat-quality measurement included three technical replicates. This batch was mainly used to evaluate relationships between type I fiber phenotypes and meat color, pH, conductivity, and intramuscular fat.

Muscle tissues were collected within 30 min after slaughter. Fiber orientation was identified during sampling to ensure that subsequent sections were perpendicular to the fiber axis. Approximately 0.8 x 0.8 x 0.8 cm3 tissue blocks were excised and placed into perforated 2.5 mL cryotubes. The perforations facilitated rapid contact between liquid nitrogen and tissue, enabling fast freezing and structural preservation. Tissue blocks were adjusted with forceps to maintain a cubic shape and avoid compression or adhesion to tube walls. Tubes were then rapidly immersed in liquid nitrogen (-196 degrees C) to minimize ice-crystal formation and preserve cross-sectional fiber structure.

### Dataset overview

See Table S4.

## METHOD DETAILS

### SDH staining and image acquisition

Frozen muscle blocks were embedded in optimal cutting temperature compound (OCT). OCT was pre-cooled at 4 degrees C to reduce bubble formation. After adding pre-cooled OCT to the embedding mold, bubbles were removed to avoid tissue-bubble contact that could lead to fiber fracture or tissue breakage during sectioning. Liquid-nitrogen-frozen muscle blocks were then placed vertically into the OCT-filled mold so that the fiber direction remained perpendicular to the cutting plane. The tissue was completely surrounded by OCT, and the mold was returned to liquid nitrogen until the OCT was fully frozen. Embedded blocks were stored at -80 degrees C.

For cryosectioning, OCT blocks containing muscle tissue were mounted on a cryostat at -20 degrees C and trimmed until a complete fiber cross-section was obtained. Serial transverse sections with a thickness of 10 micrometers were cut and attached to coated slides. Sections were required to be flat, free of obvious folds, and as perpendicular as possible to the fiber long axis.

For SDH staining, sections were placed in a humid chamber, covered with SDH staining solution, and incubated at 37 degrees C for 1-2 h. Incubation time was adjusted according to staining intensity across muscle sites. After staining and drying, warm glycerol gelatin was added near the tissue, and a coverslip was allowed to spread the mounting medium by gravity. Slides were dried in a cool ventilated area for 24 h to obtain stable stained sections for microscopy.

After staining, sections were inspected under a microscope. Images with uniform staining, intact tissue, few ice-crystal artifacts, and clear cross-sectional structure were acquired and stored as myofiber image data. The first batch yielded 235 SDH images at 1360 x 1024 resolution. The second batch yielded 222 SDH images at 1920 x 1168 resolution with matched meat-quality measurements.

### Image annotation and quality control

All SDH images underwent quality control before quantitative analysis. Inclusion criteria included intact cross-sectional tissue structure, clear staining background, identifiable type I fiber boundaries, few ice-crystal voids, and absence of obvious folding, tearing, compression artifacts, contamination, or large out-of-focus regions. Images with severe tissue damage, staining failure, excessive ice crystals, non-transverse fiber orientation, or unidentifiable type I fiber boundaries were excluded.

Initial myofiber contours were generated using Segment Anything. Candidate masks were imported into Labelme and manually corrected by trained annotators. Corrections included removal of non-fiber objects, repair of broken contours, separation of over-merged fibers, addition of missed fibers, and adjustment of ambiguous boundaries. Corrected annotations were exported as polygon coordinates and converted into binary masks and contour objects.

Quality control was performed at both image and object levels. Object-level exclusion criteria included incomplete fibers on image borders, fragments that did not represent complete cross-sections, ambiguous staining, extreme deformation, and obvious annotation errors. Final contours were required to form closed polygons. For each image, fiber-level records included polygon coordinates, contour area, centroid, bounding rectangle, perimeter, and staining intensity.

### Robust deep learning-based segmentation of skeletal muscle fibers

#### Model architecture and targeted modifications

The segmentation model was based on Mask R-CNN with a ResNet-50 feature-extraction backbone and instance mask prediction heads [45]. Five modifications were introduced to address the dominant error modes in SDH-stained myofiber images: irregular boundaries, crowded adjacent instances, small-object localization, orientation and scale variation, and computational burden.

First, modulated deformable convolution (DCNv2/mdconv) was incorporated into the ResNet-50 backbone, primarily in the conv3-conv5 stages, to enhance adaptive feature extraction for irregular fiber morphology [94]. In contrast to standard convolution with fixed sampling positions, mdconv learns both spatial offsets and modulation scalars so that the receptive field can adapt to local fiber shape:

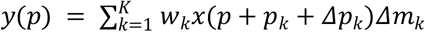

Here, *x(p)* and *y(p)* denote the input and output feature maps at position *p*, and *k* denotes the number of sampling positions. The terms indexed by *k* correspond to the sampling weight, predefined offset, learned spatial offset, and learned modulation scalar, respectively.

Second, soft-NMS replaced hard non-maximum suppression during post-processing [95]. Hard NMS can remove true neighboring fibers in crowded regions when their bounding boxes overlap. Soft-NMS instead decays detection scores according to overlap with the current highest-scoring box:

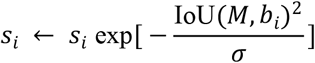

In this expression, the score term *s*_*i*_ denotes the confidence of the *i* th detection box, *M* is the current highest- scoring box, *b*_*i*_ is the ith box, and *σ* controls the strength of the overlap penalty.

Third, normalized Wasserstein distance (NWD) was incorporated into the localization objective to reduce the instability of IoU for small objects [96]. Bounding boxes were represented as Gaussian distributions, and the squared second-order Wasserstein distance between distributions *N*_*a*_ and *N*_*b*_ was written as:

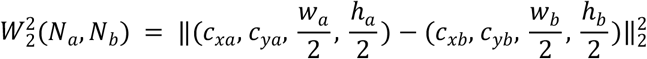

The normalized distance term was then defined as:

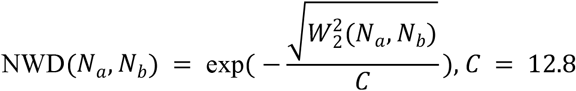

NWD provides a scale-insensitive localization term that balances loss contributions from large and small fibers.

Fourth, test-time augmentation (TTA) was used during inference to improve robustness to fiber orientation and scale. The TTA pipeline included horizontal and vertical flips, rotations, and scale variations of approximately +/-10%; predictions from augmented views were inverse-transformed and merged in the original coordinate system.

Fifth, automatic mixed precision (AMP) training was used throughout model training. The core mechanism of AMP operates by strategically managing computational precision across different operations. Specifically, it converts computationally intensive operations to FP16 precision in cases where numerical stability can be maintained, while preserving critical operations in FP32 precision where high numerical accuracy is essential. Additionally, the system maintains a master copy of weights in FP32 format while leveraging FP16 precision for gradient computation, thus establishing an optimal balance between computational efficiency and numerical stability.

The mdconv modules are primarily implemented in the backbone network, while soft-NMS is applied during the post-processing stage. AMP is utilized throughout the training process, and TTA is employed during inference. The NWD metric is incorporated into the loss function computation.

#### Training objective, inference, and post-processing

The model optimized classification, bounding-box localization, mask prediction. The total training objective was summarized as:

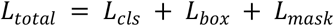

Here, the four terms denote classification loss, bounding-box regression loss, instance-mask loss, and the NWD localization term, respectively.

During inference, predictions from original and augmented views were merged after inverse coordinate transformation. Soft-NMS was then applied to retain valid crowded instances while suppressing redundant detections. Predicted masks were filtered to remove very small artifacts, small holes were filled when appropriate, obvious over-merged objects were separated when the boundary was recoverable, and accepted masks were smoothed and converted into polygon contours. These contours were passed to the same multiscale phenotype- extraction pipeline used for manual annotations.

#### Segmentation and phenotype-level evaluation

Segmentation performance was evaluated using standard object-level metrics, including average precision (AP), AP50, AP75, average recall (AR), and inference time. AP50 was defined as AP at an IoU threshold of 0.50. For each predicted and manual mask pair, overlap quality was calculated using intersection-over-union:

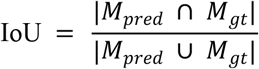

Here, the two mask terms denote the predicted and manually annotated masks, respectively. Model ablations evaluated the contribution of mdconv(Figure S15), soft-NMS, NWD, TTA, and AMP, and benchmark comparisons were performed against representative models including baseline Mask R-CNN, Cascade R-CNN, and YOLOv8.

To determine whether automated segmentation was suitable for downstream spatial phenotyping, manual and predicted contours from the same image were processed through identical feature-extraction code. Metrics included single-fiber morphology and staining features, local contact features, connected-cluster features, diffusion-ring features, and global spatial distribution features. For image i and phenotype metric k, phenotype-level relative difference was calculated as:

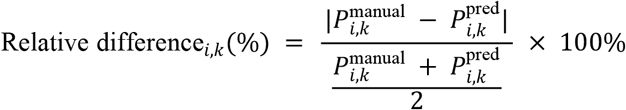

Here, the manual and predicted phenotype terms are metric values calculated from manual annotations and automated predictions, respectively. For fiber-level phenotypes, image-level summary statistics were compared; for global distribution metrics, image-level metric values were compared directly. For each model, relative differences for the same metric were averaged across all test images to obtain the cross-image mean percentage error. This phenotype-level evaluation was included because small boundary or instance-detection errors can have limited effects on AP but substantial effects on adjacency, connected-cluster assignment, and topology- derived metrics.

### Multiscale phenotype extraction of type I muscle fibers

#### Single-fiber phenotype extraction

For each segmented type I fiber ( *i*), the contour polygon was used to calculate area, perimeter, centroid, equivalent diameter, bounding rectangle, aspect ratio, circularity, rectangularity, and staining intensity.

Area *A*_*i*_ was defined as the number of pixels within the contour converted to physical area when pixel calibration was available. Equivalent diameter was calculated as:

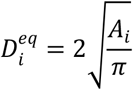

The minimum rotated rectangle of the contour yielded long and short axes *L*_*i*_ and *W*_*i*_. Aspect ratio was:

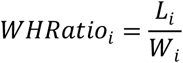

Circularity was:

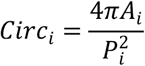

where *P*_*i*_ is perimeter. Rectangularity was:

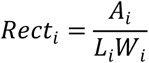

Staining intensity was calculated by creating a mask for the fiber contour and measuring average pixel intensity in RGB and grayscale channels:

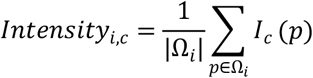

where *Ω*_*i*_ is the set of pixels inside the mask of fiber (i), and *I*_*c*_(*p*) is the intensity of pixel *p* in channel *c*.

#### Spatial adjacency and connected-cluster analysis

To quantify local contact between type I fibers, we first calculated the minimum contour distance between any two fibers:

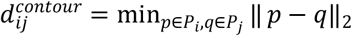

Centroid distance was calculated as:

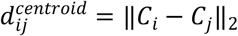

An adjacency matrix (*Adj*) was constructed from contour distances. Two fibers were considered adjacent when their minimum contour distance was less than half the minimum short-axis length among fibers in the image:

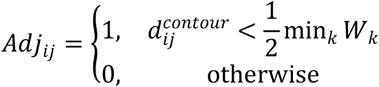

where *W*_*k*_ is the short-axis length of the minimum bounding rectangle of fiber (*k*). Myofibers were treated as graph nodes *V* and adjacency relationships as edges *E*, yielding a type I fiber adjacency graph (*G* = (*V, E*)). Connected components were identified by breadth-first search and defined as connected clusters.

For connected cluster (*c*), with node set *V*_*c*_ and edge set *E*_*c*_, the within-cluster fiber count was:

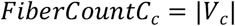

Cluster area was:

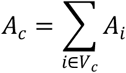

Cluster contours were generated by merging contours of fibers in the cluster. Based on the merged contour, cluster aspect ratio, circularity, rectangularity, and compactness were calculated. If 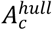 is the merged contour or hull area, compactness was defined as:

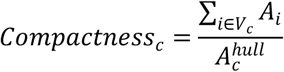

Cluster circularity was:

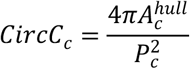

where ( *p*_*c*_ ) is the perimeter of the merged contour. Within-cluster connection density was described by the average clustering coefficient. For node (*i*), its degree (*k*_*i*_), and number of edges among its neighbors (*e*_*i*_), the local clustering coefficient was:

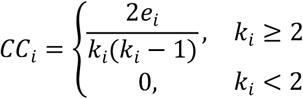

The cluster-level coefficient was:

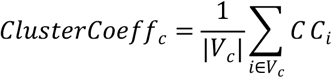

Graph centrality metrics including degree centrality, closeness centrality, betweenness centrality, and eigenvector centrality were calculated for individual fibers within connected clusters. OutRatio and InRatio were used to describe direct contact strength. For fiber (*i*), contour points with distances below contact threshold (*τ*) to neighboring fibers were treated as contact-participating boundary points. OutRatio describes the proportion of the current fiber boundary involved in contact, whereas InRatio summarizes the contact proportion contributed by neighboring fibers toward the current fiber.

#### Diffusion-ring microenvironment analysis

To characterize the local microenvironment centered on each type I fiber, we constructed a diffusion-ring region. For fiber (*i*), the circle centered at centroid ( *C*_*i*_ ) used a radius equal to 2.5 times the mean equivalent diameter 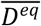 of type I fibers in the image:

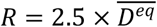

The diffusion disk was:

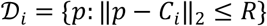

The set of fibers falling within this disk was:

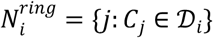

The local type I fiber area fraction, 2.5AED-TMCR, was:

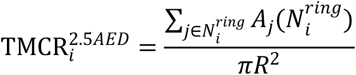

where 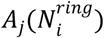 is the intersection area of the fiber *j* and the diffusion disk. Relative coordinates of neighboring fibers within the diffusion disk were:

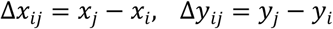

For diffusion-ring clustering, each neighborhood was encoded as a binary image in which pixels of type I fiber regions were assigned 1 and background 0. Similarity between two diffusion disks (*a*) and (*b*) was:

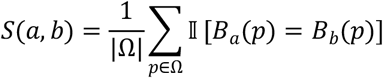

K-means clustering was then performed using the similarity or corresponding distance matrix, and within- class mean similarity was used to evaluate consistency of microenvironmental spatial patterns.

#### Global spatial phenotype calculation

Global image-level phenotypes summarized type I fiber abundance, spatial density, clustering, dispersion, and complexity. Ratio of Stained Cells was calculated as the total type I fiber area divided by image area:

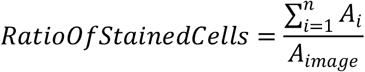

Global packing density was calculated as the total type I fiber area divided by the convex hull area of all type I fibers:

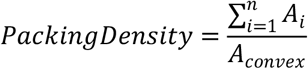

The nearest-neighbor index (NNI) was used to quantify global aggregation or dispersion of type I fiber centroids. If (*r*_*i*_) is the distance from each centroid to its nearest neighbor, the observed mean nearest-neighbor distance was:

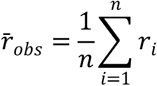

The expected nearest-neighbor distance under spatial randomness was:

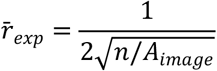

NNI was:

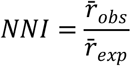

Under this definition, NNI < 1 generally indicates more clustering than random expectation, whereas NNI > 1 indicates dispersion.

Global distance entropy and angle entropy were used to describe spatial complexity. Pairwise centroid distances or angles were binned equidistantly to obtain bin probabilities (*p*_*b*_), and entropy was:

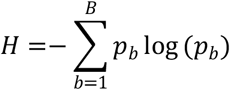

where *B* is the number of bins. Higher entropy indicates more complex or heterogeneous distance or angular distribution of type I fibers.

#### Topology Enhancement Score, TES

To distinguish type I fiber abundance from spatial topology enhancement, we constructed the Topology Enhancement Score (TES). The rationale is that many spatial topology metrics depend on the overall proportion of SDH-positive type I fibers. Raw topology metrics therefore cannot be directly interpreted as independent topology organization capacity. To obtain abundance-adjusted topology, Ratio of Stained Cells was used as a covariate. Direction coefficients of TES implementation seeTable S5

For sample (*s*) and topology metric (*m*), the within-breed model was:

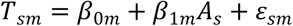

where (*T*_*sm*_) is topology metric (*m*) in sample (*s*), (*A*_*s*_) is Ratio of Stained Cells, and (*ε*_*sm*_) is the residual topology deviation after removing the abundance effect. For a pooled cross-breed model, breed can be included as an additional covariate:

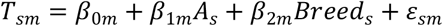

The residual was defined as the abundance-adjusted topology residual. A positive residual indicates that the observed topology is higher than expected given the type I fiber proportion, whereas a negative residual indicates lower-than-expected topology.

Because different topology metrics have different biological directions, residuals were direction-corrected before TES integration. For metrics where higher values indicate stronger clustering, connectivity, local enrichment, or organization, the direction coefficient was set to (*d*_*m*_). For metrics where higher values indicate dispersion, greater inter-cluster distance, or weaker organization, the direction coefficient was set to (d_m_ =− 1). Direction-corrected residuals were then Z-score standardized:

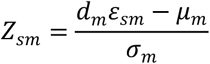

where (*µ*_*m*_) and (*σ*_*m*_) are the mean and standard deviation of the direction-corrected residuals for metric (*m*). TES was defined as:

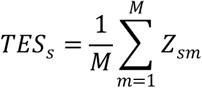

where (*M*) is the number of topology metrics included. Thus, TES > 0 indicates stronger-than-expected spatial organization at a given type I fiber abundance, whereas TES < 0 indicates weaker-than-expected organization.

#### Spatial transcriptomics integration

To evaluate the relationship between type I myofiber topology and local molecular programs, spatial transcriptomics data from PMM, SM, and related skeletal muscle sections were integrated with type I myofiber instance segmentations from matched histological images. Normalized expression matrices were obtained from h5ad files. Spot expression values were taken from spot_adata.X, registered spatial coordinates were taken from deconv_adata.obsm[“spatial_reference”], and spots were matched between AnnData objects using shared obs_names. Gene-expression matrices, spot coordinates, tissue images, and myofiber polygons were transformed into a common histological reference-image coordinate system.

Each spatial transcriptomic spot was represented as a circular region centered at its registered coordinate, using the corresponding reference_spot_radius or spot_radius. The overlap ratio between spot s and fiber f was calculated as:

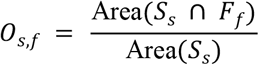

where O_s,f_ is the overlap ratio between spatial transcriptomic spot s and fiber f, S_s_ is the circular region representing spot s, F_f_ is the polygon region of fiber f, and Area(·) denotes the area of the corresponding region. This ratio was used to assign spot-level molecular information to segmented myofibers.

A spot was considered matched to a fiber when the overlap ratio exceeded the preset threshold (0.4) used for the analysis. Fibers without valid matched spots were excluded from downstream molecular analyses. For fibers matched to multiple spots, all matched spot indices, overlap ratios, and expression matrices were retained; for correlation analyses and visualization, the expression profile of the spot with the largest overlap ratio was used as the representative molecular signal to reduce dilution from neighboring fibers.

After spot-fiber matching, expression profiles were merged with single-fiber phenotype data to create a fiber-level object containing gene expression, spatial coordinates, contour information, diffusion-ring metrics, graph centrality metrics, and connected-cluster identifiers. Connected clusters containing fewer than five fibers were excluded from cluster-level molecular analyses. The final fiber-level dataset included topology metrics such as OutRatio, InRatio, 2.5AED-TMCR, degree centrality, betweenness centrality, closeness centrality, eigenvector centrality, and clustering coefficient.

Target genes included MYH7, TNNT1, TNNC1, TPM3, CRYAB, CSRP3, ANKRD2, and RRAD. GAPDH, ACTB, and B2M were included as control genes. A slow-oxidative module was defined using MYH7, TNNT1, TNNC1, and TPM3; a stress/cytoskeletal module was defined using CRYAB, CSRP3, ANKRD2, and RRAD. For each module m, expression values of genes in the module were Z-score standardized across fibers and averaged to obtain a module score:

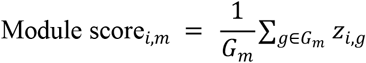

where Module score_i,m_ is the score of module m in fiber i, G_m_ is the set of genes included in module m, |G_m_| is the number of valid genes in the module, and z_i,g_ is the Z-score-standardized expression of gene g in fiber i.

Global Moran’s I was used to test whether target genes and module scores showed non-random spatial organization. A k-nearest-neighbor spatial weight matrix was constructed from fiber centroid coordinates, with k = 8 in the PMM sample, and was row-normalized before analysis. For expression vector *x*, Global Moran’s I was calculated as:

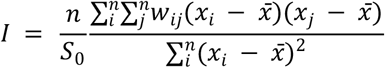

where I is Global Moran’s I, n is the number of matched fibers, w_ij_ is the spatial weight between fibers i and j, S_0_ is the sum of all spatial weights, x_i_ and x_j_ are expression or module-score values for fibers i and j, and 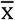 is the mean expression or module-score value across fibers.

Statistical significance was assessed using 999 permutations. Local Moran’s I was then calculated under the same spatial weight matrix to identify fiber-level local expression domains. Local P values were corrected using the Benjamini-Hochberg false discovery rate procedure. Fibers were assigned to high-high, low-low, high-low, or low-high Moran quadrants based on centered expression and spatial lag. High-high hotspots were defined as fibers whose expression and neighborhood-average expression were both above the global mean and significant after FDR correction.

Molecular hotspots were linked to type I myofiber topology using three complementary analyses. First, topology metrics were compared between fibers inside and outside molecular high-high hotspots. High-topology fibers were defined using the median value of each topology metric, enrichment in molecular hotspots was tested using a hypergeometric test, and hotspot versus non-hotspot differences were evaluated using the Mann-Whitney U test. Second, molecular high-high hotspots were compared with phenotype high-high domains for OutRatio, InRatio, 2.5AED-TMCR, and centrality-related metrics. Spatial overlap was quantified using the Jaccard index:

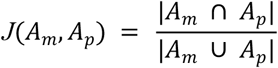

where J is the Jaccard index, A_m_ is the set of fibers in a molecular high-high hotspot, and A_p_ is the set of fibers in a phenotype high-high domain. The numerator is the overlap between the two sets, and the denominator is their union.

Permutation tests with 999 random hotspot-label shuffles were used to generate null distributions for the Jaccard index, followed by FDR correction. Third, Spearman correlations were calculated between fiber-level Local Moran’s I values and topology metrics across all gene/module-phenotype combinations, with FDR correction. In connected-cluster-level analyses, gene expression or module scores were averaged across fibers within each cluster. Clusters above the 70_th_ percentile of clustering coefficient were defined as the high-ClusterCoeff group, whereas clusters below the 30_th_ percentile were defined as the low-ClusterCoeff group. Differences between these groups were tested using Welch’s t test and the Mann-Whitney U test, and Cohen’s d was used to estimate effect size.

#### Meat quality measurement and phenotype association

The second sample batch was used to evaluate associations between type I myofiber phenotypes and meat- quality traits. Traits included meat color, pH, and conductivity measured at 45-60 min and 24 h postmortem, as well as intramuscular fat content (IMF) in longissimus dorsi muscle (LDM) samples. Each meat-quality measurement included three technical replicates, and replicate means were used for downstream analyses.

Meat color was measured using a CR-400 colorimeter after removing surface fascia and fat. The muscle cross-section was placed flat, and the colorimeter lens was kept perpendicular to the tissue surface with full contact. The aperture diameter was 8 mm, illuminant C was used, and the white calibration plate was set to 100. L*, a*, and b* values were measured at 45-60 min and 24 h postmortem. pH was measured using a carcass meat- quality pH meter after probe rinsing and calibration. The probe was inserted into muscle while avoiding fat and connective tissue. Conductivity was measured using a carcass meat-quality conductivity meter calibrated with a 0 S/m block, and readings were recorded after stabilization. IMF was measured by petroleum ether Soxhlet extraction.

Image-derived phenotype matrices were matched to meat-quality records using breed, individual ID, muscle type, and replicate ID parsed from image names. When multiple images were available for the same animal and muscle type, phenotype metrics were averaged before merging with meat-quality measurements. Analyses of meat color, pH, and conductivity included LDM and psoas major muscle (PMM), whereas IMF analyses were restricted to LDM samples. Only individuals with both valid image-derived phenotypes and meat-quality measurements were retained for the corresponding analysis.

The phenotype matrix included single-fiber morphology, staining intensity, global spatial distribution, diffusion-ring microenvironment, and connected-cluster topology metrics. Representative metrics included Area, MaxD, WHRatio, Rect, Circ, Redness, Greenness, Blueness, Grayness, Ratio of Stained Cells, Packing Density, NNI, Cluster Distance, Number of Clusters, Global ED, Global EA, OutRatio, InRatio, 2.5AED-TMCR, clustering coefficient, FiberCount-C, WHRatio-C, and Rect-C. Metrics were organized into individual, distribution, interaction, and cluster feature groups and merged by individual ID. Missing values after merging were imputed using the feature mean.

Single-variable associations between image-derived phenotypes and meat-quality traits were evaluated using Spearman rank correlation:

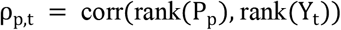

where ρ_p,t_ is the Spearman correlation coefficient between phenotype metric p and meat-quality trait t, P_p_ is the vector of image-derived phenotype values for metric p, Y_t_ is the vector of trait values for trait t, and rank(·) denotes rank transformation before correlation.

Combinations with fewer than three valid observations were not tested. For LDM IMF, Spearman correlations were calculated separately between each image-derived phenotype and the mean IMF value.

Partial least squares regression (PLSR) was used to evaluate the joint explanatory capacity of multivariate type I myofiber phenotypes. For each meat-quality trait, a PLS1 model was built with the trait as the single response variable and the type I myofiber phenotype matrix as predictors. Predictors and response variables were Z-score standardized before modeling:

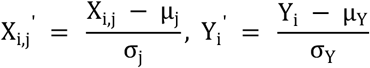

where X_i,j_ is the value of predictor j in sample i, Y_i_ is the response value in sample i, μ_j_ and σ_j_ are the mean and standard deviation of predictor j, and μ_Y_ and σ_Y_ are the mean and standard deviation of the response variable. 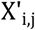 and 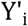 denote standardized predictor and response values used for PLS modeling.

The PLS1 model was written as:

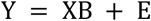

where Y is the n by 1 standardized response vector for one meat-quality trait, X is the n by p standardized predictor matrix of type I myofiber phenotype metrics, B is the p by 1 vector of PLS regression coefficients, E is the residual vector, n is the number of samples, and p is the number of predictors.

The number of PLS components was selected by five-fold cross-validation. Candidate component numbers were evaluated by prediction residual sum of squares (PRESS), and the component number that minimized PRESS was selected:

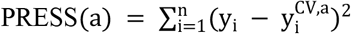

where PRESS(a) is the prediction residual sum of squares obtained using a PLS components, y_i_ is the observed response value for sample i, y^CV^_i,a_ is the cross-validated prediction for sample i from the model with a components, and n is the number of samples. The component number with the lowest PRESS was selected.

The final model was refitted using the selected number of components. Regression coefficients, component loadings, predicted values, and variable importance in projection (VIP) scores were then calculated. For predictor j, VIP was calculated as:

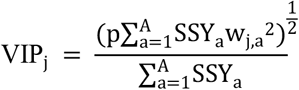

where VIP_j_ is the variable importance in projection score for predictor j, p is the total number of predictors, A is the number of retained PLS components, SSY_a_ is the amount of response-variable variation explained by component a, and w_j,a_ is the normalized weight of predictor j on component a. A VIP > 1 threshold was used to identify predictors with high contribution to the PLS model.

Predictors with VIP > 1 were treated as candidate phenotype variables with high contribution to the corresponding trait. Model performance was evaluated using the coefficient of determination (R2) and mean absolute relative error (MARE):

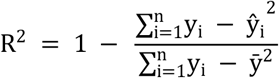

where R^2^ is the coefficient of determination, y_i_ is the observed trait value for sample i, ŷ_i_ is the modelpredicted value, 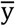 is the mean observed trait value, and n is the number of samples. R^2^ measures the proportion of trait variation explained by the model.

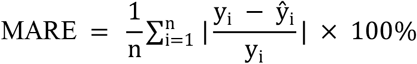

where MARE is the mean absolute relative error, y_i_ is the observed trait value, ŷ_i_ is the predicted trait value, and n is the number of samples. For conventional meat-quality traits, MARE was calculated after transforming predictions back to the original trait scale.

For conventional meat-quality traits, predictions were transformed back to the original measurement scale before calculating R2 and MARE. To assess the added contribution of spatial topology, control models were constructed after removing spatial-distribution, local-adjacency, and connected-cluster metrics, including NNI, Cluster Distance, Number of Clusters, Global ED, Global EA, OutRatio, InRatio, 2.5AED-TMCR, clustering coefficient, FiberCount-C, WHRatio-C, and Rect-C. Control models used the same PLS1 workflow, normalization, cross-validation, PRESS-based component selection, VIP calculation, and performance evaluation. IMF was analyzed separately using LDM samples, with the mean IMF value as the response variable and matched LDM image-derived phenotypes as predictors.

## QUANTIFICATION AND STATISTICAL ANALYSIS

Unless otherwise specified, statistical analyses were performed in R and Python. Continuous variables were analyzed at the level of independent images, muscle sites, or animals, depending on the corresponding experimental question. Variables were summarized using the mean, median, standard deviation, and interquartile range as appropriate. Before multivariate analyses, phenotype metrics with different units were standardized column-wise:

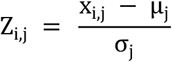

where Z_i,j_ is the standardized value of metric j in sample i, x_i,j_ is the original value, and μ_j_ and σ_j_ are the mean and standard deviation of metric j across samples. This standardization places metrics with different units on a common scale before multivariate analysis.

Two-group comparisons were performed using the Wilcoxon rank-sum test, also referred to as the Mann-Whitney U test, unless a different test was specified for the analysis. For analyses requiring parametric comparison of means, Welch’s t test was used. Multi-group comparisons were performed using the Kruskal-Wallis test or ANOVA according to the distribution and analysis design. Effect size for two-group comparisons was estimated using Cohen’s d:

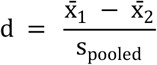

where d is Cohen’s d, 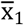 and 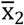 are the group means, and s_pooled_ is the pooled standard deviation. Cohen’s d was used to summarize the direction and magnitude of two-group differences in addition to P values.

Correlations among phenotype metrics, between spatial transcriptomic module scores and type I myofiber topology, and between meat-quality traits and image-derived phenotypes were evaluated using Spearman rank correlation. Correlation coefficients are reported as Spearman’s ρ.

Multivariate phenotype structure was analyzed using principal component analysis (PCA), uniform manifold approximation and projection (UMAP), K-means clustering, and hierarchical clustering. PCA was used to identify major axes of variation in the high-dimensional phenotype matrix. UMAP was applied to PCA-reduced components to visualize nonlinear sample structure. K-means clustering was used to identify muscle sites or image groups with similar spatial phenotype patterns. Hierarchical clustering was performed on Z-score- standardized phenotype matrices using Euclidean distance and Ward.D2 linkage and was used to visualize similarity among muscle sites, breeds, or phenotype modules.

Global differences among breeds, anatomical regions, or phenotype classes in multivariate phenotype space were tested using permutational multivariate analysis of variance (PERMANOVA) based on Euclidean distance matrices with 999 permutations. PERMANOVA was applied separately to phenotype sets such as spatial- distribution metrics, morphology metrics, connected-cluster metrics, staining-intensity metrics, and diffusionring microenvironment metrics to evaluate whether grouping factors explained overall multivariate variation.

For analyses involving multiple comparisons, including correlation screening, local spatial autocorrelation in spatial transcriptomic analyses, and multimetric group comparisons, P values were corrected using the Benjamini-Hochberg false discovery rate procedure. Unless otherwise stated, nominal significance was defined as P < 0.05, and multiple-testing-adjusted significance was defined as FDR < 0.05. Significance in figures was annotated as *P < 0.05, **P < 0.01, and ***P < 0.001.

### Association of Muscle Spatial Topology with Bulk Transcriptomic Profiles

This study used spatial phenotype samples as anchors to precisely match muscle spatial measurements with pig BodyMap bulk RNA-seq data[97]. The expression data were obtained from GSE162145_pig_PCG.all_exp_tpm_gene.txt, with expression values reported as TPM. The expression matrix was organized with genes as rows and samples as columns. The spatial phenotype variables included ClusterCoefft, FiberCount-C, NNI, ClusterDistance, PackingDensity, Ratio of Stained Cells, OutRatio, and 2.5AED-TMCR. The spatial phenotype measurements were precisely matched to the BodyMap bulk expression data at the sample level. In total, 98 spatial phenotype-bulk transcriptome pairs covering 39 muscle sites were obtained. Samples from different muscle sites were not averaged in advance; each successfully matched bulk expression profile was treated as an independent observation in the subsequent models.

Based on the sample-pairing results, the expression matrix was extracted and reordered according to bulk_sample_id. Lowly expressed genes were filtered by retaining only those with TPM > 1 in at least 20% of the samples. The expression values were then log-transformed as follows:

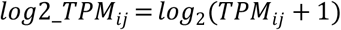

where *i* denotes a gene and *j* denotes a sample. The filtered and transformed expression matrix was used for topology-score construction, single-gene modeling, predefined transcriptional module scoring, and genomewide ranking analysis.

Next, PCA and PLS models were trained using the RC pig meat-quality phenotype data and the corresponding muscle spatial measurements, yielding fixed PCA parameters, PLS weights, standardization parameters, orientation parameters, and score-scaling parameters. All of these parameters were derived from the RC pig meat-quality data and were not retrained using the current 98 bulk-matched samples. First, LocalDensityCompositeScore was constructed using the fixed PCA parameters for NNI, PackingDensity, and 2.5AED-TMCR. Subsequently, each meat-quality trait was used to construct an independent trait-specific topology-score based on five spatial inputs: ClusterCoefft, OutRatio, ClusterDistance, FiberCount-C, and LocalDensityCompositeScore. NNI, PackingDensity, and 2.5AED-TMCR were not used as independent PLS inputs; instead, they were first integrated into LocalDensityCompositeScore. The topology-score was calculated as follows:

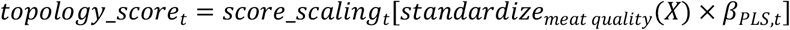

where *X* denotes the five spatial input variables, *β*_*PLS,t*_ denotes the fixed PLS weights trained for trait *t* using the RC pig meat-quality data, *standardize*_*meat quality*_ denotes standardization using the mean and standard deviation of the meat-quality data, and *score_scaling*_*t*_ denotes the fixed score-scaling parameter for trait *t* Separate topology-scores were constructed for each meat-quality trait, and no cross-trait averaged topology-score was calculated.

After obtaining the trait-specific topology-scores, single-gene association analyses were performed separately for each trait. The 39 exact muscle sites were included as no-intercept fixed effects, and the trait- specific topology-score and standardized type I fiber proportion were included as explanatory variables:

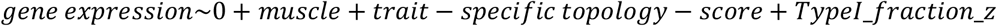

The models were fitted using the lmFit function in limma, followed by robust empirical Bayes variance estimation using eBayes(trend = TRUE, robust = TRUE). For each gene, the effect estimate, moderated *t* - statistic, and *p*-value were obtained. BH FDR correction was performed within each trait, followed by global BH FDR correction across traits.

Building on the single-gene analyses, nine predefined transcriptional modules were subsequently analyzed. The modules were predefined according to muscle biology and established biological knowledge. Their names and predefined gene compositions are provided in **Table S6**. For each module, the expression of each module gene was first standardized across all samples. The mean standardized expression of the successfully matched genes within each sample was then calculated as the module score:

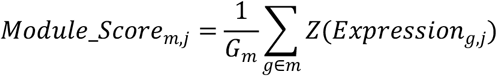

where *m* denotes a module, *j* denotes a sample, *G*_*m*_ denotes the number of successfully matched genes in module *m*, and *Z*(*Expression*_*g,j*_) denotes the standardized expression of gene *g* in sample *j* . The module-level model used the same fixed effects and covariates as the gene-expression model:

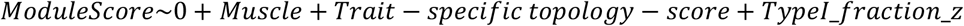

For each trait, BH multiple-testing correction was first performed across the nine modules, followed by global BH FDR correction across all trait-module combinations.

To complement the predefined module analysis, genome-wide gene rankings were generated using the moderated *t*-statistics obtained from the gene-expression model for each trait. Pig gene symbols were mapped to Entrez IDs using org.Ss.eg.db. When multiple gene symbols mapped to the same Entrez ID, the record with the largest absolute *t*-statistic was retained. GO Biological Process enrichment analysis was then performed using clusterProfiler. GO gene sets were restricted to sizes ranging from 10 to 500 genes, and 10,000 simple permutations with a fixed random seed were used for each trait. The resulting enrichment statistics were corrected using trait-specific BH FDR and global BH FDR across traits. GO terms with substantial hierarchical overlap, such as RNA processing, mRNA processing, and RNA splicing, were interpreted as shared functional themes rather than as independent findings.

After the primary model, predefined module analysis, and GSEA were completed, muscle-classification sensitivity analyses were performed to assess the sensitivity of module effects to the definition of muscle background. The primary analysis used the 39 exact muscle sites as fixed effects. The sensitivity analyses separately replaced the exact muscle-site terms with developmental_context, replaced them with detailed_origin_compartment, or retained exact muscle-site adjustment within the major_meat_subset. The specific classifications and corresponding muscle sites are provided in **Table S7**. The major meat-muscle subset included BB, ISP, SSP, SUB, TBLO, TBM, TM, LD, LDM, PP, PS, ADD, BF, GM, GRA, PM, RF, SAR, SM, ST, VI, and VM. This subset was used only for sensitivity analysis. All three alternative models retained the trait- specific topology-score and Ratio of Stained Cells. Subsequently, the nine module effects for the same trait from each sensitivity model were paired with the corresponding effects from the primary model. Pearson correlation coefficients and effect-direction concordance rates were calculated to evaluate consistency across model specifications. This analysis was used to assess model stability, and the different muscle-classification models were not treated as independent experimental replicates.

Finally, to distinguish the effects of the composite topology-score from those of individual spatial measurements, each of the seven raw spatial metrics was standardized and independently included in separate models. This analysis did not use meat-quality traits, PLS weights, LocalDensityCompositeScore, or trait-specific topology-scores. The gene-expression model for the raw spatial metrics was:

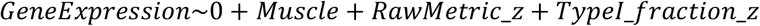

The corresponding module-level model was:

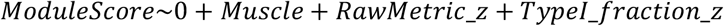

Each raw spatial metric was modeled separately, and BH FDR correction was performed within the corresponding gene-level or module-level comparison set. This analysis was used to determine whether the observed transcriptional associations depended on a single spatial metric or required the integration of multiple spatial dimensions.

## Notes

### Competing Interest Statement

The authors have declared no competing interest.

