## Supplemental files for "A spatial topology atlas of type I myofibers in porcine skeletal muscle reveals local slow-oxidative niches associated with meat quality traits"

**Table S1. Overview of the multiscale type I myofiber spatial phenotype system.**

| **Phenotype layer** | **Metric class** | **Metric** | **Metric-level interpretation** |
| --- | --- | --- | --- |
| Single-fiber attributes | Size | **Area** | Cross-sectional area of each SDH-positive type I myofiber; reflects fiber size and local hypertrophy or atrophy. |
| Single-fiber attributes | Size | **MaxD** | Maximum fiber diameter or long-axis length derived from the contour or bounding rectangle; captures fiber caliber and elongation scale. |
| Single-fiber attributes | Morphology | **WHRatio** | Width-to-height or long-to-short axis ratio of the fitted rectangle; describes fiber elongation and shape anisotropy. |
| Single-fiber attributes | Morphology | **Rect** | Rectangularity of the fiber contour; measures how closely the fiber fills its minimum bounding rectangle. |
| Single-fiber attributes | Morphology | **Circ** | Circularity of the fiber contour; describes roundness and boundary regularity of individual type I fibers. |
| Single-fiber attributes | Staining intensity | **Redness / Greenness / Blueness** | Mean RGB channel intensities within the fiber mask; capture color-channel variation in SDH-stained fibers. |
| Single-fiber attributes | Staining intensity | **Grayness** | Mean grayscale intensity within the fiber mask; provides an image-level proxy for SDH staining intensity and oxidative phenotype. |
| Local adjacency | Direct contact | **OutRatio** | Fraction of the current fiber boundary participating in contact with neighboring type I fibers; quantifies outward same-type contact strength. |
| Local adjacency | Direct contact | **InRatio** | Contact contribution from neighboring type I fibers toward the current fiber; quantifies local inward contact pressure or neighborhood contact intensity. |
| Local adjacency | Graph centrality | **DegCent** | Degree centrality of a fiber in the adjacency graph; measures how many same-type neighbors are directly connected to the fiber. |
| Local adjacency | Graph centrality | **BetCent** | Betweenness centrality; identifies fibers that bridge shortest paths within a connected type I fiber cluster. |
| Local adjacency | Graph centrality | **CloCent** | Closeness centrality; measures how readily a fiber can reach other fibers within the same connected cluster. |
| Local adjacency | Graph centrality | **EigCent** | Eigenvector centrality; assigns higher values to fibers connected to other highly connected fibers, reflecting network influence. |
| Connected-cluster structure | Cluster size | **FiberCount-C** | Number of type I fibers within a connected cluster; reflects the scale of local slow-oxidative structural units. |
| Connected-cluster structure | Cluster size | **Area-C** | Merged area of all type I fibers in a connected cluster; summarizes cluster-level tissue occupancy. |
| Connected-cluster structure | Cluster connectivity | **ClusterCoeff** | Average clustering coefficient within a connected cluster; quantifies how tightly neighboring fibers interconnect. |
| Connected-cluster structure | Cluster shape | **WHRatio-C** | Aspect ratio of the merged cluster contour; describes elongation of the connected type I fiber unit. |
| Connected-cluster structure | Cluster shape | **Rect-C** | Rectangularity of the merged cluster contour; captures cluster shape regularity relative to its bounding rectangle. |
| Connected-cluster structure | Cluster shape | **Circ-C** | Circularity of the merged cluster contour; indicates whether the connected cluster is compact and rounded or irregular. |
| Connected-cluster structure | Cluster compactness | **Compactness-C** | Ratio of merged fiber area to enclosing hull or contour area; measures how densely type I fibers occupy the cluster domain. |
| Diffusion-ring microenvironment | Local enrichment | **2.5AED-TMCR** | Type I fiber area fraction within a disk centered on each fiber with radius 2.5 times the mean equivalent diameter; measures local same-type enrichment. |
| Diffusion-ring microenvironment | Local abundance | **Fiber count in 2.5AED disk** | Number of type I fibers located within the diffusion-ring neighborhood; captures local microenvironmental cellular density. |
| Diffusion-ring microenvironment | Spatial displacement | **Relative neighbor coordinates** | Coordinates of neighboring fibers relative to the central fiber; describe directional displacement and local spatial arrangement. |
| Diffusion-ring microenvironment | Neighborhood pattern | **Diffusion-ring similarity / class** | Similarity or K-means class of binary diffusion-ring patterns; summarizes recurring local spatial microenvironment types. |
| Global tissue configuration | Abundance | **Ratio of Stained Cells** | Total SDH-positive type I fiber area divided by image area; measures tissue-level type I fiber abundance. |
| Global tissue configuration | Compactness | **Packing Density** | Total type I fiber area divided by the convex hull area of all type I fibers; quantifies overall compactness of type I fiber distribution. |
| Global tissue configuration | Aggregation / dispersion | **NNI** | Nearest-neighbor index of type I fiber centroids; values below 1 indicate clustering, whereas values above 1 indicate dispersion. |
| Global tissue configuration | Cluster abundance | **Number of Clusters** | Number of connected components in the type I fiber adjacency graph; reflects whether type I fibers form many separated units or fewer integrated structures. |
| Global tissue configuration | Cluster spacing | **Cluster Distance** | Mean distance among connected-cluster centroids; captures separation among type I fiber clusters at the tissue scale. |
| Global tissue configuration | Spatial complexity | **Global ED** | Entropy of pairwise centroid-distance distribution; measures distance heterogeneity and global spatial complexity. |
| Global tissue configuration | Spatial complexity | **Global EA** | Entropy of pairwise centroid-angle distribution; measures directional heterogeneity of type I fiber spatial arrangement. |

***Table S2. Key resources.***

| **Resource** | **Source** | **Identifier / version** |
| --- | --- | --- |
| Rongchang pigs | This paper | Female, ~10 months, ~75 kg |
| Large White pigs | This paper | Female, ~10 months, ~153 kg |
| Landrace pigs | This paper | Female, ~10 months, ~153 kg |
| SDH staining reagents | This paper | Succinate dehydrogenase staining |
| Segment Anything Model | Meta AI | SAM2.1 |
| Labelme | Labelme | labelme 5.8.1 |
| Python | Python Software Foundation | Python 3.11.15 |
| R | R Foundation | R-4.5.1 |
| PyTorch / Detectron2 / MMDetection | PyTorch / Detectron2 / MMDetection | Deep learning training framework |
| OpenCV | OpenCV | Contour and geometric feature extraction |
| libpysal / esda | libpysal / esda | libpysal 4.14.1;esda 2.9.0. Spatial transcriptomics and spatial autocorrelation analysis |

***Table S3. Skeletal muscle abbreviations and English full names used in the manuscript.***

| **Body region** | **Abbreviation** | **Full name** |
| --- | --- | --- |
| Forelimb skeletal muscle | BB | Biceps brachii |
| Forelimb skeletal muscle | DAP | Deltoid, acromial part |
| Forelimb skeletal muscle | DSP | Deltoid, scapular part |
| Forelimb skeletal muscle | EDC | Common digital extensor |
| Forelimb skeletal muscle | EDLF | Lateral digital extensor (forelimb) |
| Forelimb skeletal muscle | FDPF | Deep digital flexor (forelimb) |
| Forelimb skeletal muscle | ISP | Infraspinatus |
| Forelimb skeletal muscle | SSP | Supraspinatus |
| Forelimb skeletal muscle | SUB | Subscapularis |
| Forelimb skeletal muscle | TBLA | Triceps brachii, lateral head |
| Forelimb skeletal muscle | TBLO | Triceps brachii, long head |
| Forelimb skeletal muscle | TBM | Triceps brachii, medial head |
| Forelimb skeletal muscle | TM | Teres major |
| Trunk skeletal muscle | BRA | Brachiocephalicus |
| Trunk skeletal muscle | Dia | Diaphragm |
| Trunk skeletal muscle | LD | Latissimus dorsi |
| Trunk skeletal muscle | LDM | Longissimus dorsi |
| Trunk skeletal muscle | OEA | External abdominal oblique |
| Trunk skeletal muscle | PP | Deep pectoral |
| Trunk skeletal muscle | PS | Superficial pectoral |
| Trunk skeletal muscle | RA | Rectus abdominis |
| Trunk skeletal muscle | RHO | Rhomboideus |
| Trunk skeletal muscle | SP | Splenius |
| Trunk skeletal muscle | TA | Transversus abdominis |
| Trunk skeletal muscle | TRC | Trapezius, cervical part |
| Hindlimb skeletal muscle | ADD | Adductor |
| Hindlimb skeletal muscle | BF | Biceps femoris |
| Hindlimb skeletal muscle | EDLH | Lateral digital extensor (hindlimb) |
| Hindlimb skeletal muscle | FDPH | Deep digital flexor (hindlimb) |
| Hindlimb skeletal muscle | FDSH | Superficial digital flexor (hindlimb) |
| Hindlimb skeletal muscle | GASL | Gastrocnemius, lateral head |
| Hindlimb skeletal muscle | GASM | Gastrocnemius, medial head |
| Hindlimb skeletal muscle | GM | Gluteus medius |
| Hindlimb skeletal muscle | GRA | Gracilis |
| Hindlimb skeletal muscle | PMM | Psoas major |
| Hindlimb skeletal muscle | Pop | Popliteus |
| Hindlimb skeletal muscle | RF | Rectus femoris |
| Hindlimb skeletal muscle | SAR | Sartorius |
| Hindlimb skeletal muscle | SM | Semimembranosus |
| Hindlimb skeletal muscle | SOL | Soleus |
| Hindlimb skeletal muscle | ST | Semitendinosus |
| Hindlimb skeletal muscle | TFL | Tensor fasciae latae |
| Hindlimb skeletal muscle | TIC | Tibialis cranialis |
| Hindlimb skeletal muscle | VI | Vastus intermedius |
| Hindlimb skeletal muscle | VL | Vastus lateralis |
| Hindlimb skeletal muscle | VM | Vastus medialis |

**Table S4. Dataset overview.**

| **Dataset** | **Breed** | **Region / muscle site** | **Animals** | **Images** | **Meat-quality records** | **Resolution** | **Main use** |
| --- | --- | --- | --- | --- | --- | --- | --- |
| Dataset 1 (first batch) | RC + LW | All sampled sites (46 muscle sites) | 4 RC + 4 LW | 233 | NA | 1360 x 1024 | Multi-site and multi-breed type I fiber atlas; topology enhancement score (TES); neighbor-dependent clustering. |
| Dataset 1 (first batch) | Rongchang pig (RC) | All sampled sites (42 muscle sites) | 4 RC | 102 | NA | 1360 x 1024 | Breed-level annotation summary for topology phenotyping. |
| Dataset 1 (first batch) | Rongchang pig (RC) | Front limbs | 4 RC | 31 | NA | 1360 x 1024 | Regional SDH-positive type I fiber distribution. |
| Dataset 1 (first batch) | Rongchang pig (RC) | Trunk | 4 RC | 31 | NA | 1360 x 1024 | Regional SDH-positive type I fiber distribution. |
| Dataset 1 (first batch) | Rongchang pig (RC) | Hind limbs | 4 RC | 40 | NA | 1360 x 1024 | Regional SDH-positive type I fiber distribution. |
| Dataset 1 (first batch) | Large White pig (LW) | All sampled sites (42 muscle sites) | 4 LW | 131 | NA | 1360 x 1024 | Breed-level annotation summary for topology phenotyping. |
| Dataset 1 (first batch) | Large White pig (LW) | Front limbs | 4 LW | 35 | NA | 1360 x 1024 | Regional SDH-positive type I fiber distribution. |
| Dataset 1 (first batch) | Large White pig (LW) | Trunk | 4 LW | 35 | NA | 1360 x 1024 | Regional SDH-positive type I fiber distribution. |
| Dataset 1 (first batch) | Large White pig (LW) | Hind limbs | 4 LW | 61 | NA | 1360 x 1024 | Regional SDH-positive type I fiber distribution. |
| Dataset 2 (second batch) | LW + Landrace (LR) | PMM, LDM (2 muscle sites) | 20 LW + 17 LR | 222 | 74 | 1920 x 1168 | Type I fiber atlas; meat-quality association; partial least squares (PLS) modeling; spatial metric contribution. |
| Dataset 2 (second batch) | Large White pig (LW) | PMM, LDM (2 muscle sites) | 20 LW | 120 | 40 | 1920 x 1168 | Breed-specific subset with matched meat-quality measurements. |
| Dataset 2 (second batch) | Landrace pig (LR) | PMM, LDM (2 muscle sites) | 17 LR | 102 | 34 | 1920 x 1168 | Breed-specific subset with matched meat-quality measurements. |
| Spatial transcriptomics | Landrace pig(PMM),Large White pig (SM) | PMM,SM (2 muscle sites) | 1LR+1LW | 2 | NA | 2000x1868 | Coupling between molecular programs and spatial topology. |
| bulk RNA-seq Data | Rongchang pig (RC) | All sampled sites (42 muscle sites) | 4 RC | 102 | NA | 1360 x 1024 | Bulk Spatial Topology and Muscle Transcriptome Association Analysis |

***Table S5 Direction coefficients of TES implementation.***

| **Metric group** | **Candidate metrics** | **Direction for TES** | **Note** |
| --- | --- | --- | --- |
| Local contact | OutRatio, InRatio | + | Higher values indicate stronger local contact |
| Connected-cluster topology | ClusterCoeff, FiberCount-C | + | Higher values indicate stronger within-cluster connectivity or larger cluster size |
| Local enrichment | 2.5AED-TMCR | + | Higher values indicate stronger local type I fiber enrichment |
| Compactness | Packing Density | + | Higher values indicate more compact global distribution |
| Distance/dispersion | Cluster Distance | - | Larger distances indicate more separated clusters and should be reversed |
| Cluster number | Number of Clusters | - | At fixed abundance, more clusters may indicate dispersion or multicenter organization |
| Nearest-neighbor index | NNI | - | Under the standard definition, NNI < 1 indicates clustering, whereas NNI > 1 indicates dispersion. |

***Table S6. Predefined transcriptional modules and constituent genes***

| **Module** | **Predefined genes** |
| --- | --- |
| slow contractile | MYH7, TNNC1, TNNI1, MYL2, ACTN2, CSRP3 |
| oxidative mitochondrial | PPARGC1A, COX4I1, COX5B, NDUFS1, ATP5F1A, SDHA, TFAM |
| fatty acid oxidation | CPT1B, ACADM, ACADVL, HADHA, HADHB, PPARA, FABP3 |
| fast glycolysis | PFKM, ALDOA, ENO3, GAPDH, PGAM2, PKM, LDHA, MYH1 |
| vascular capillary | VEGFA, KDR, VWF, PECAM1, EMCN, RAMP2 |
| ECM remodeling | COL1A1, COL3A1, COL5A1, DCN, LUM, MMP2, TIMP1 |
| calcium excitation | RYR1, CACNA1S, ATP2A1, ATP2A2, CASQ1, TRDN |
| lipid deposition | PPARG, CEBPA, FABP4, PLIN1, ADIPOQ, DGAT2 |
| immune inflammation | TYROBP, LST1, CSF1R, CD74, HLA-DRA, C1QA |

***Table S7. Muscle-site classifications used for primary and sensitivity analyses***

| **Classification variable** | **Category** | **Included muscle sites** |
| --- | --- | --- |
| developmental_context | axial_body_wall_somitic | LDM, SP, OEA, RA, TA |
| developmental_context | forelimb_pectoral_hypaxial | BB, DAP, DSP, EDC, EDLF, FDSH, ISP, SSP, SUB, TBLO, TBM, TM, BRA, LD, PP, PS, RHO, TRC |
| developmental_context | hindlimb_hypaxial | ADD, BF, GASL, GASM, GM, GRA, PM, RF, SAR, SM, SOL, ST, TIC, VI, VL, VM |
| detailed_origin_compartment | epaxial_axial | LDM, SP |
| detailed_origin_compartment | hypaxial_body_wall_respiratory | OEA, RA, TA |
| detailed_origin_compartment | cervico_scapular_pectoral_transitional | BRA, LD, PP, PS, RHO, TRC |
| detailed_origin_compartment | forelimb_proper | BB, DAP, DSP, EDC, EDLF, FDSH, ISP, SSP, SUB, TBLO, TBM, TM |
| detailed_origin_compartment | hindlimb_proper | ADD, BF, GASL, GASM, GM, GRA, PM, RF, SAR, SM, SOL, ST, TIC, VI, VL, VM |

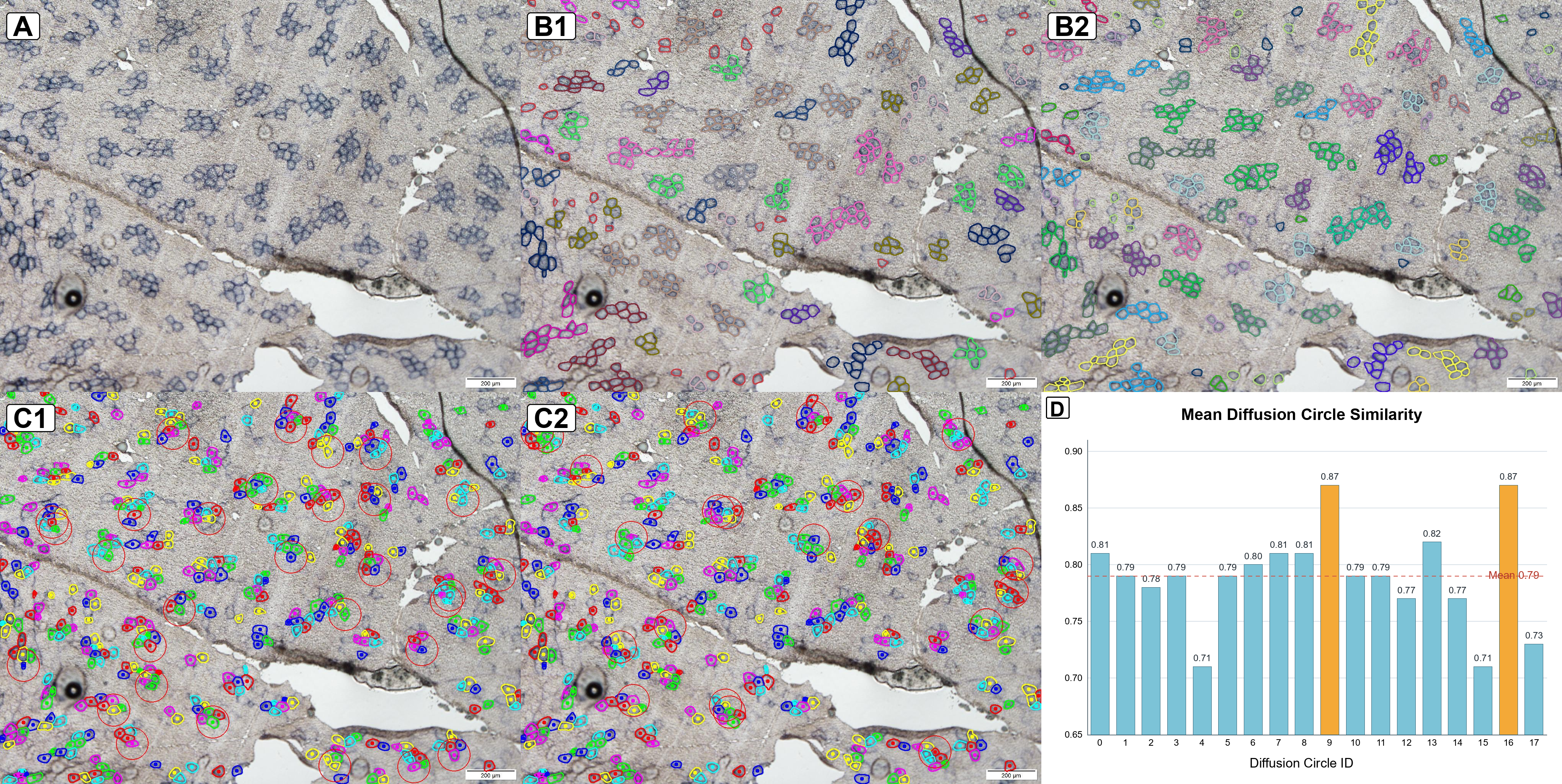
**Figure S1. SDH-stained image, myofiber-contour annotation, and local microenvironment classification.** A shows a representative original image of ADD. B1 shows connected clusters of different classes obtained using the weighted K-means method, and B2 shows the results obtained using the PCA-K-means method. C1 and C2 show two classes of diffusion rings clustered by the K-means method. D shows the intra-ring similarity levels for each category.

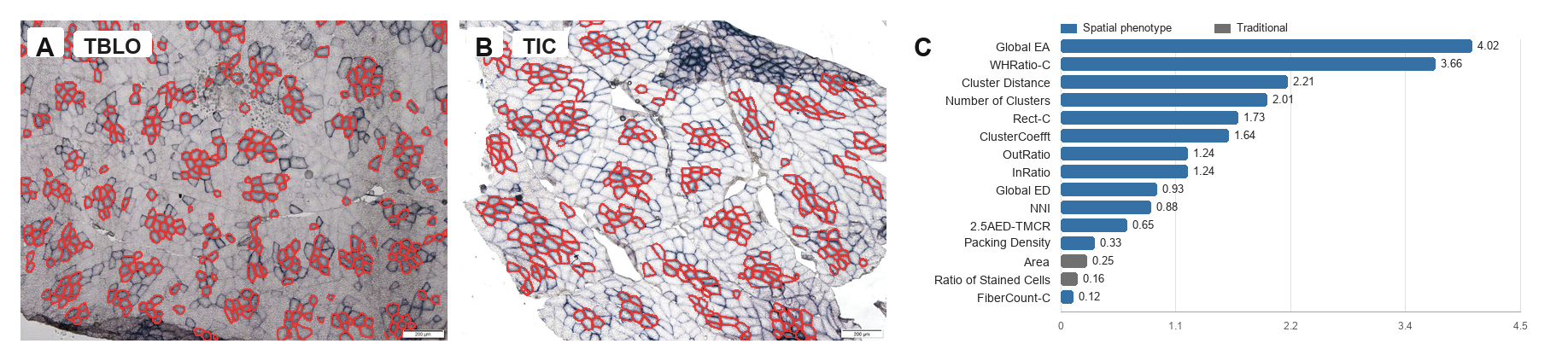

***Figure S2. Spatial phenotype differences beyond conventional proportion metrics. Representative muscle comparisons show that similar type I fiber proportions can correspond to different cluster numbers, inter-cluster distances, adjacency patterns, and global spatial complexity.*** *(A, B) Muscle sections from TBLO and TIC within the large-white (LW) breed. (C) Absolute differences in phenotype metrics between TBLO and TIC, sorted from largest to smallest.*

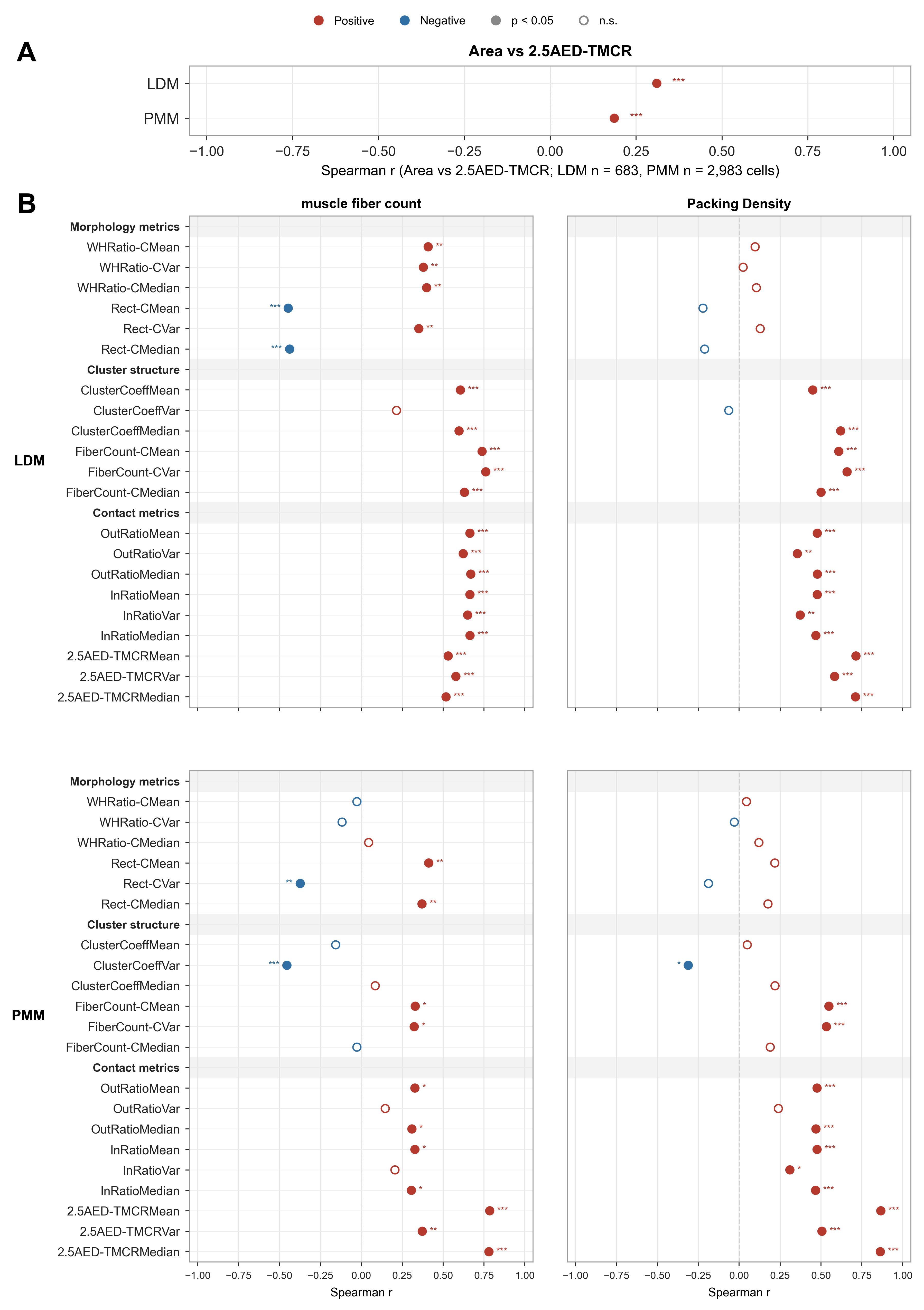
***Figure S3. Relationship between single-fiber phenotype and local topological position in LW pigs.Type I fibers with stronger local contact or higher centrality tended to show larger cross-sectional areas or altered spatial organization.*** *(A)Muscle-site-level pooled correlations between type I fiber cross-sectional area (Area) and the 2.5AED-TMCR (ratio of the total fiber area to its fitted circle, 2.5×). All fibers of a muscle site were pooled into a single Spearman correlation (LDM, n = 683 fibers from 4 images; PMM, n = 2,983 fibers from 4 images); the number of pooled fibers is given under the x axis, and the correlation magnitude is read from the position along the x axis. (B)Site-specific validation in the longissimus dorsi (LDM) and psoas major (PMM), showing correlations between the two image-level phenotypes (muscle fiber count and packing density) and type I fiber morphology, cluster-structure and contact metrics. Red and blue indicate positive and negative correlations, respectively; filled bubbles indicate significant correlations. In both panels circles follow the same encoding, and the legend at the top of the figure is shared by panels A and B: red and blue denote positive and negative correlations, respectively, filled circles mark significant correlations (p < 0.05)* *and open circles non-significant ones (n.s.), with *p < 0.05, **p < 0.01, ***p < 0.001. PMM samples are the PM image files, reported as PMM throughout.*

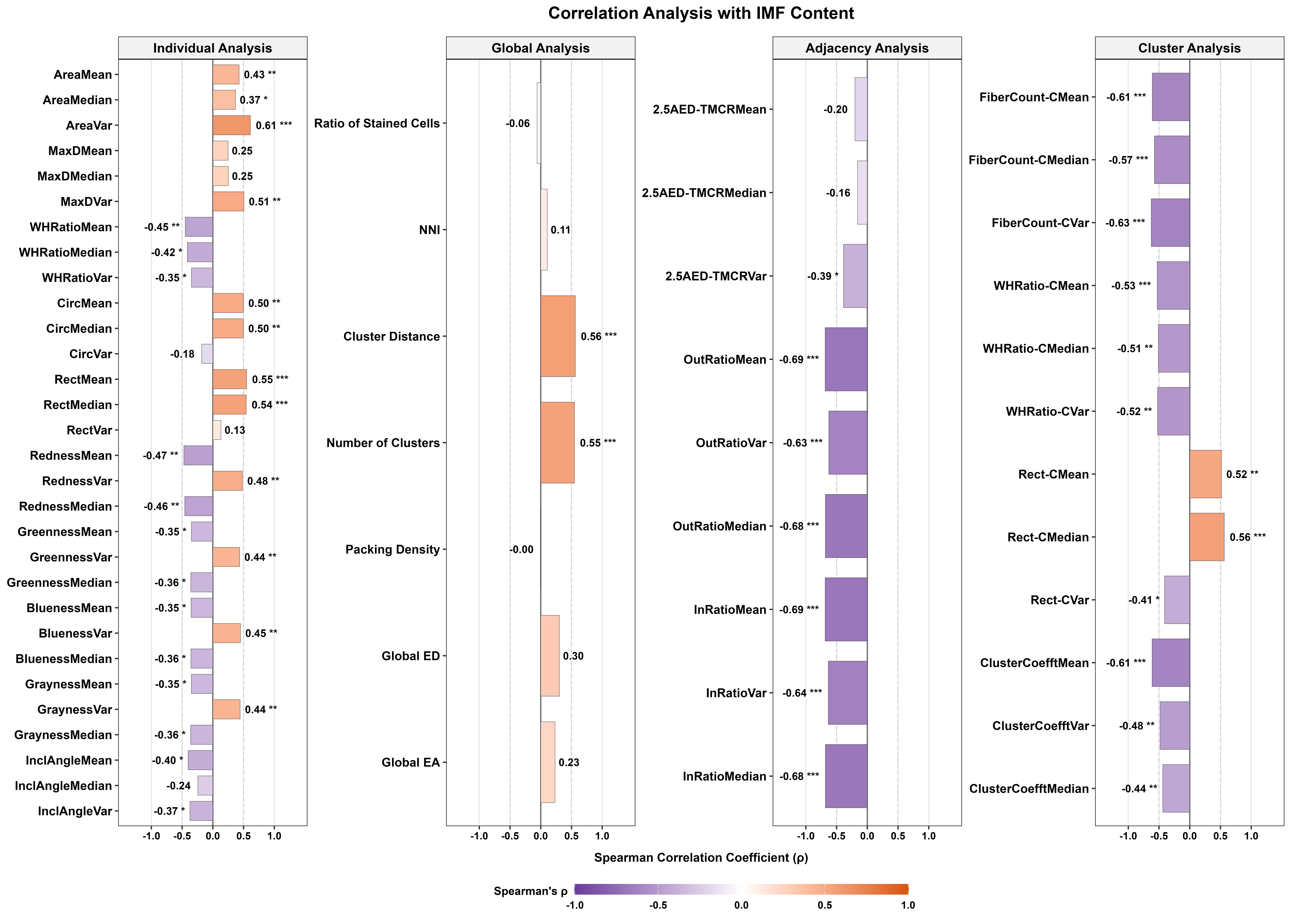

**Figure S4. Correlation matrix between intramuscular fat (IMF) and type I fiber phenotypes.**

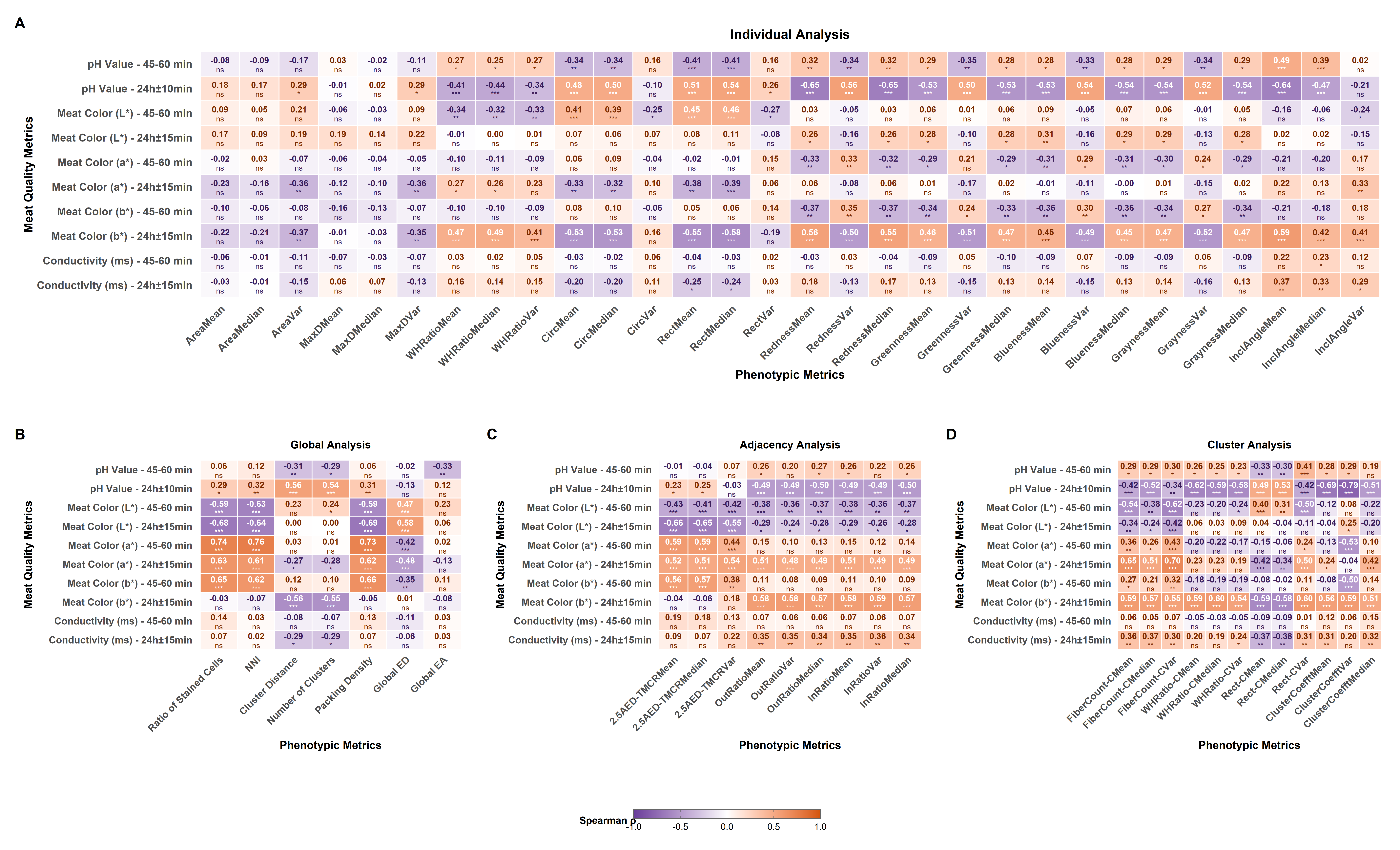
***Figure S5. Correlation matrix between other meat-quality traits and type I fiber phenotypes.***

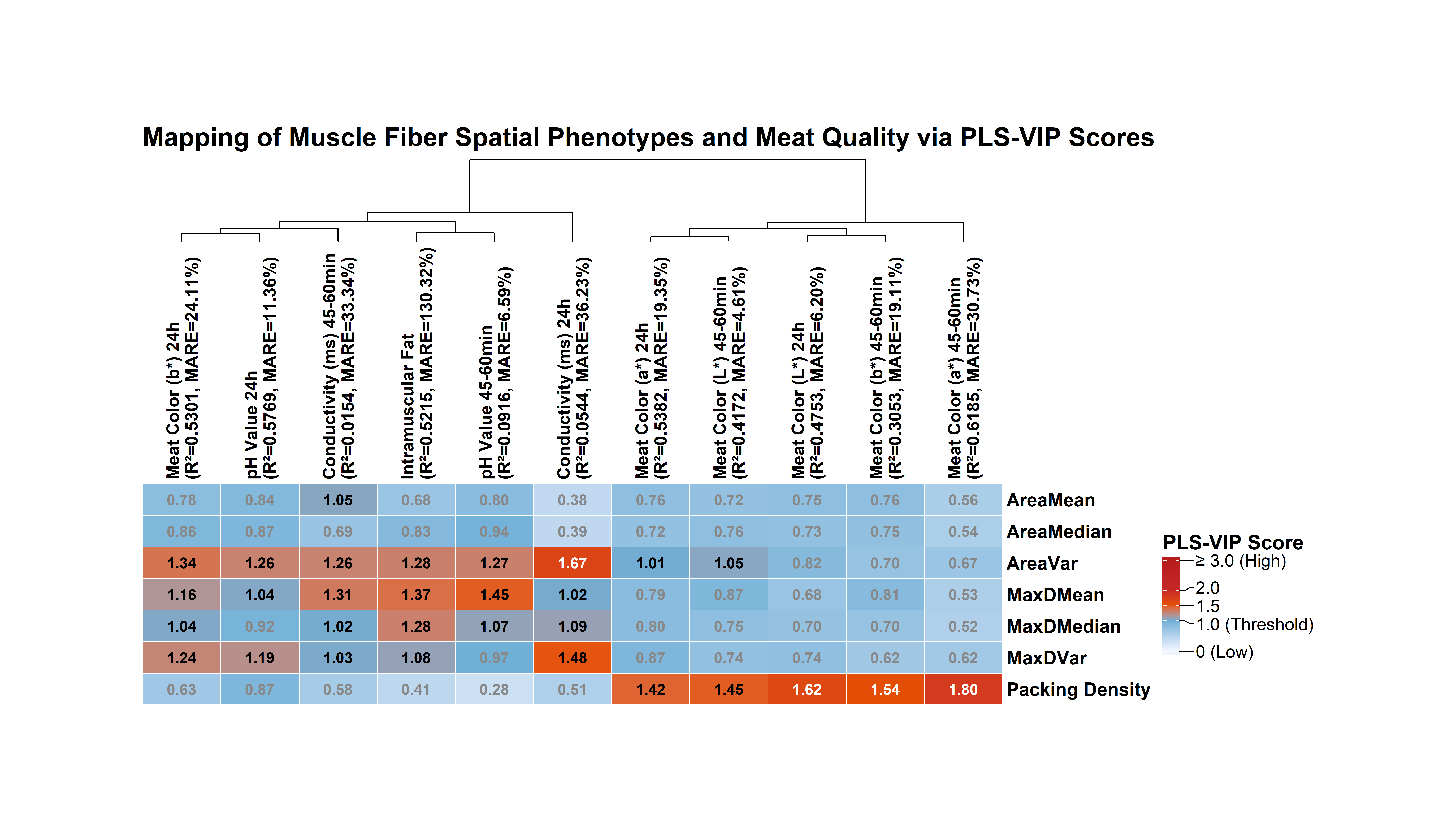

**Figure S6. Control models excluding spatial metrics show that topology provides information beyond conventional proportion and morphology for explaining meat quality.**

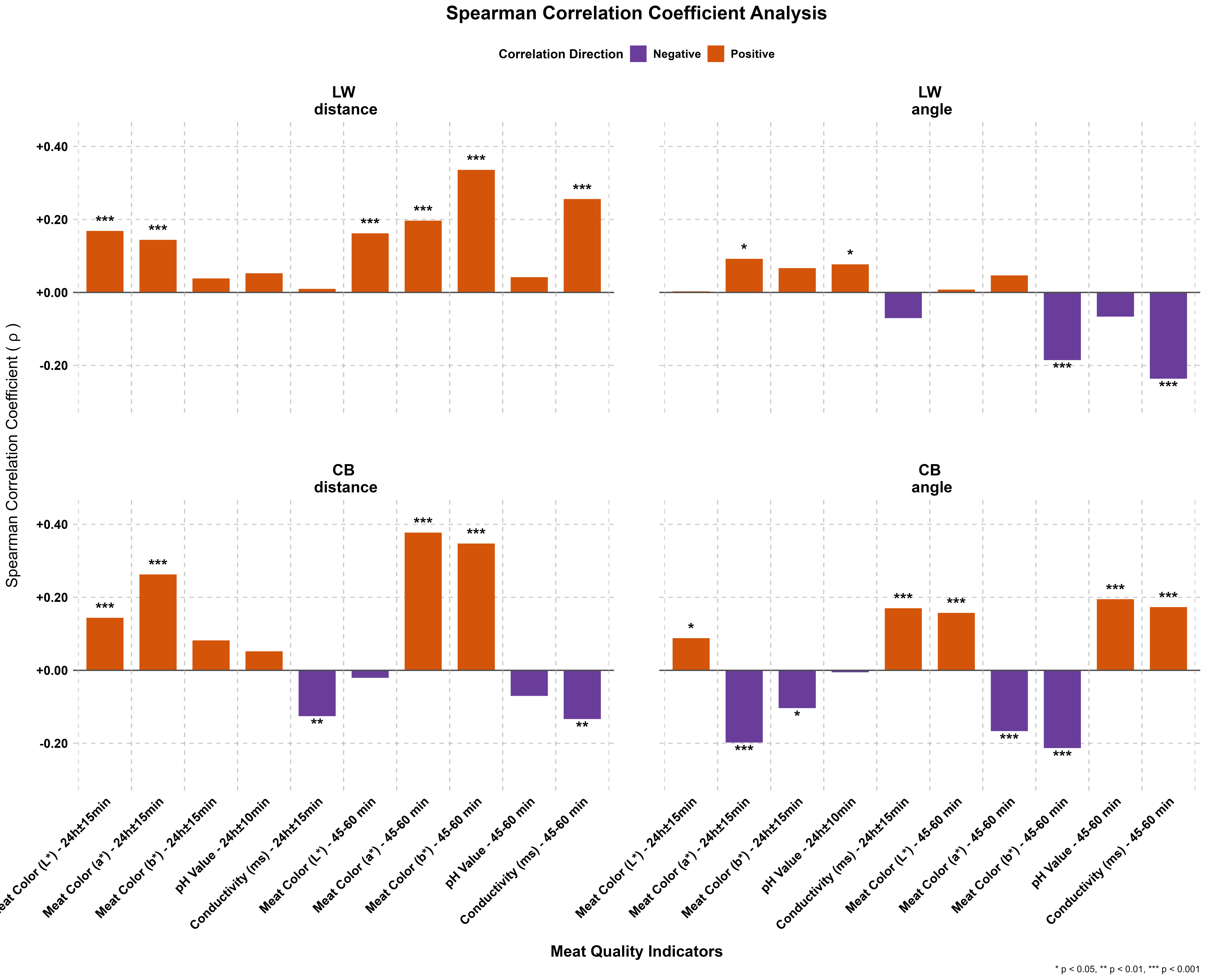

**Figure S7. Correlation analysis between meat-quality differences and the differences of global spatial distribution fitting based on Gaussian radial basis functions.**

***Figure S8. Adjusted associations between meat-quality trait-specific topology scores and prespecified transcriptional modules.*** *The heatmap shows 99 adjusted regression effects between 11 meat-quality trait-specific topology scores and 9 prespecified transcriptional modules. Models were adjusted for exact muscle site and Type I fiber fraction. Red and blue indicate positive and negative effects, respectively; the cell values indicate the magnitude of the regression coefficients. All models included 98 exact phenotype–bulk RNA-seq matches spanning 39 muscle sites. The heatmap displays the direction and magnitude of effects and does not encode statistical significance. None of the associations survived trait-wise or global FDR correction.*
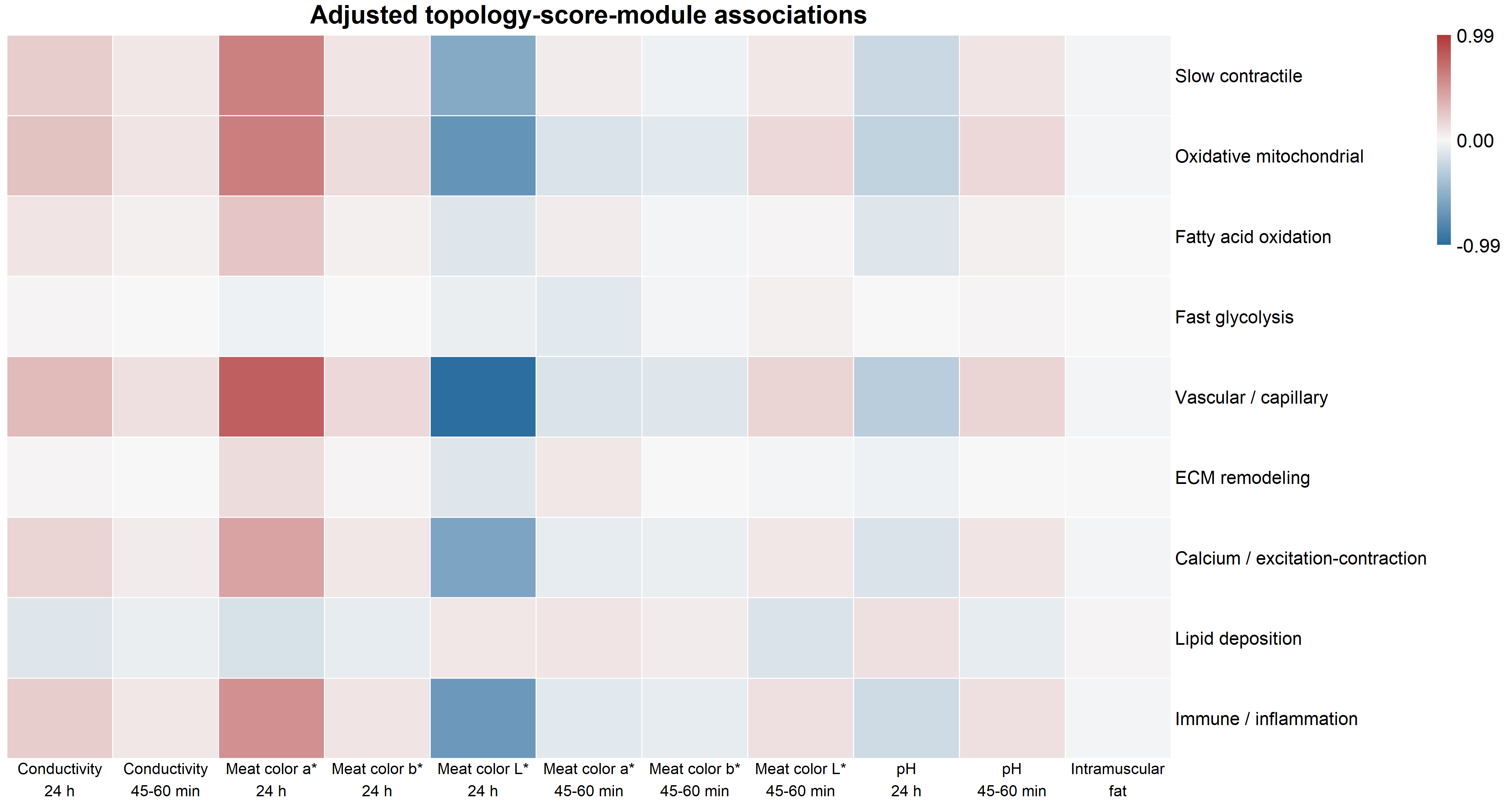

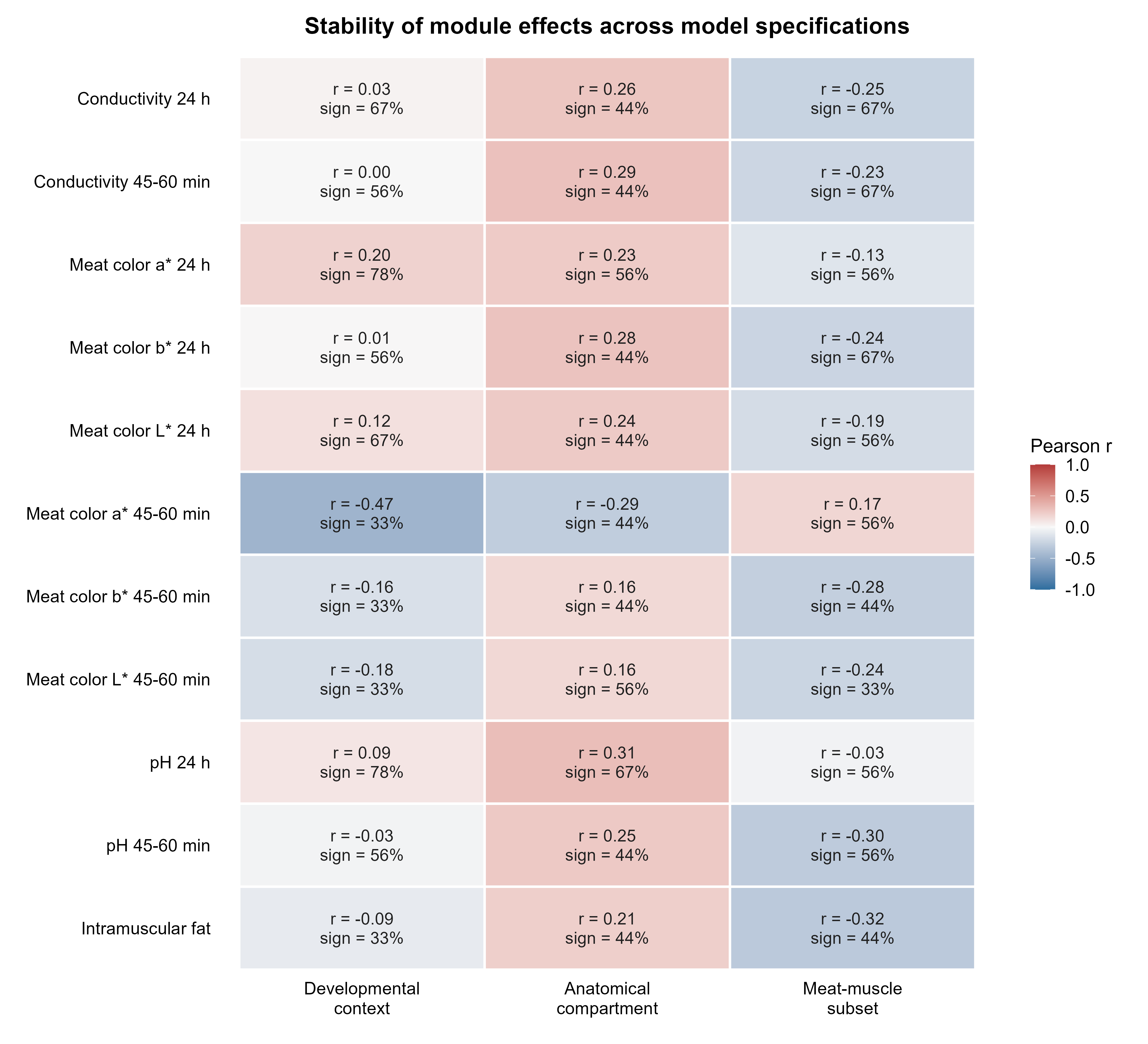
***Figure S9. Stability of module effects across muscle classifications and sample subsets.*** *This figure compares module effects from developmental-origin-adjusted, anatomical-compartment-adjusted, and major meat-muscle subset models with those from the primary exact-muscle-site model. Each cell is calculated from the same 9 prespecified modules for a given meat-quality trait. Colour indicates the Pearson correlation between the alternative-model and primary-model effects; each cell also reports the correlation coefficient and the proportion of effects with concordant signs. Because each correlation is based on only 9 modules, this figure is intended to describe model sensitivity and does not constitute independent statistical validation. Overall, module effects were sensitive to muscle classification and sample selection.*

***Figure S10. Adjusted associations between raw spatial metrics and prespecified transcriptional modules.*** *The heatmap shows 63 independent adjusted models testing each of 7 raw spatial metrics against each of 9 prespecified transcriptional modules. Each model used one raw spatial metric standardized across the 98 matched bulk samples and was adjusted for exact muscle site and Type I fiber fraction. These models did not use PLS weights, meat-quality trait information, or composite topology scores. Red and blue indicate positive and negative effects, respectively; the values displayed within cells are nominal P values, whereas the colour scale represents the magnitude of the regression coefficients. None of the combinations reached nominal P < 0.05 or FDR < 0.05.*
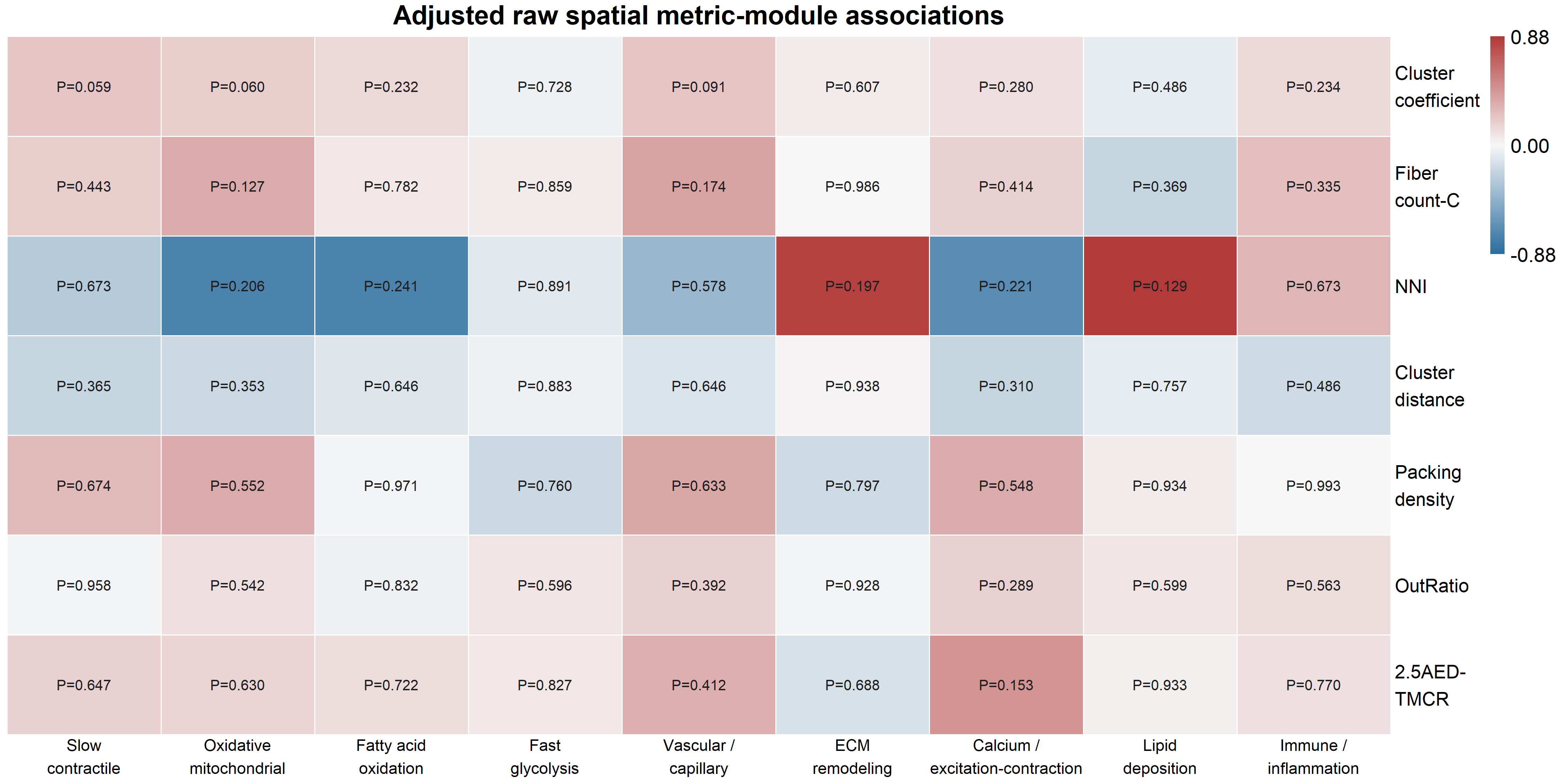

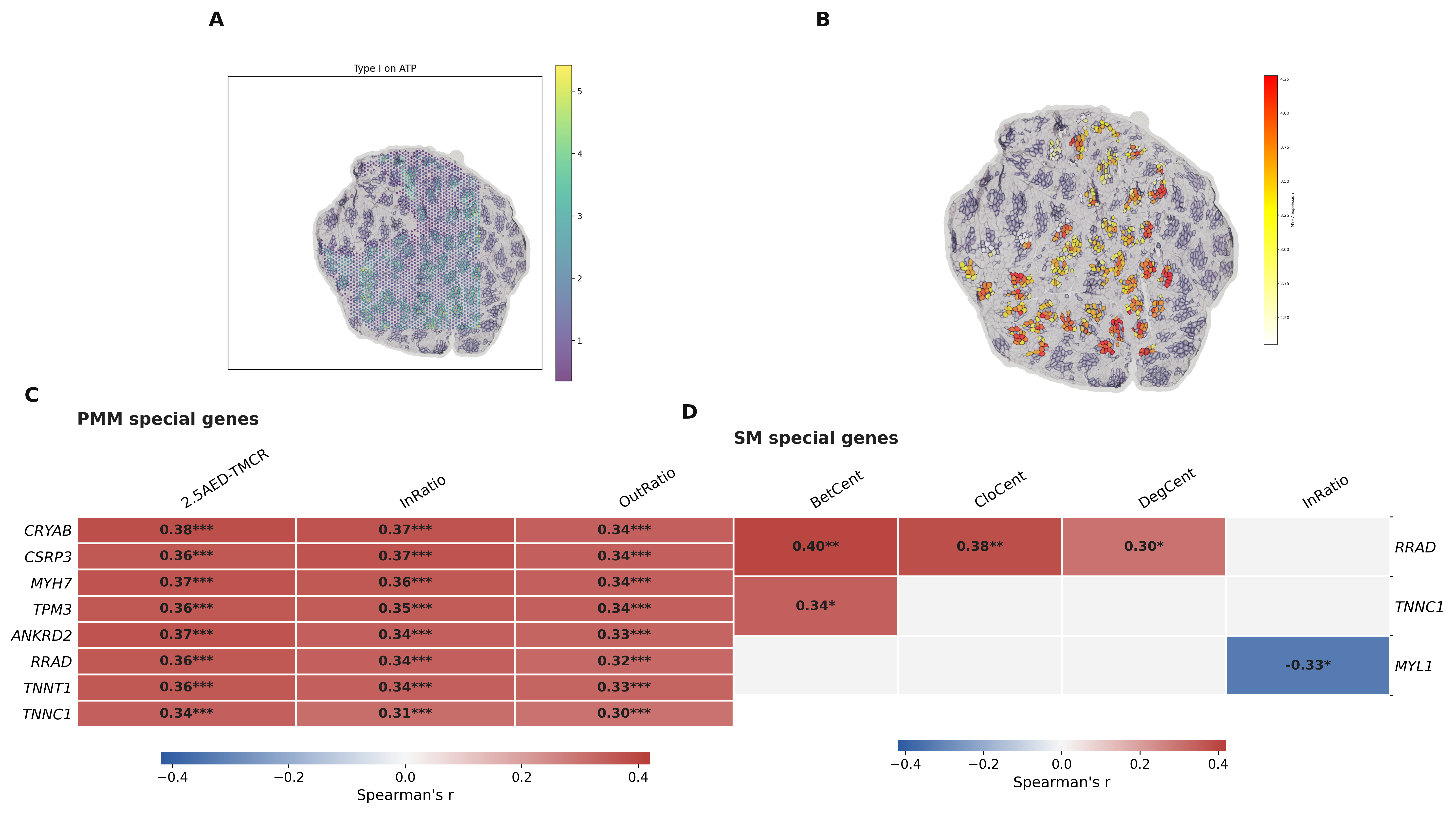

**Figure S11. Gene-phenotype correlation analysis linking spatial transcriptomics to type I fiber topology.** Multiple genes related to slow-oxidative metabolism, mechanical adaptation, and metabolic homeostasis are correlated with local adjacency or centrality metrics in PMM and SM. (A) Distribution of PMM alignment sites (B) Expression of MYH7 gene in PMM, in some regions, the number of muscle fibers is too low because some cluster sites are sparse, or some muscle fibers at the junctions do not have effective sites, resulting in cluster breakage and being filtered out (C) Correlation coefficients between some genes and phenotypes in PMM. (D)Correlation coefficients between some genes and phenotypes in SM.

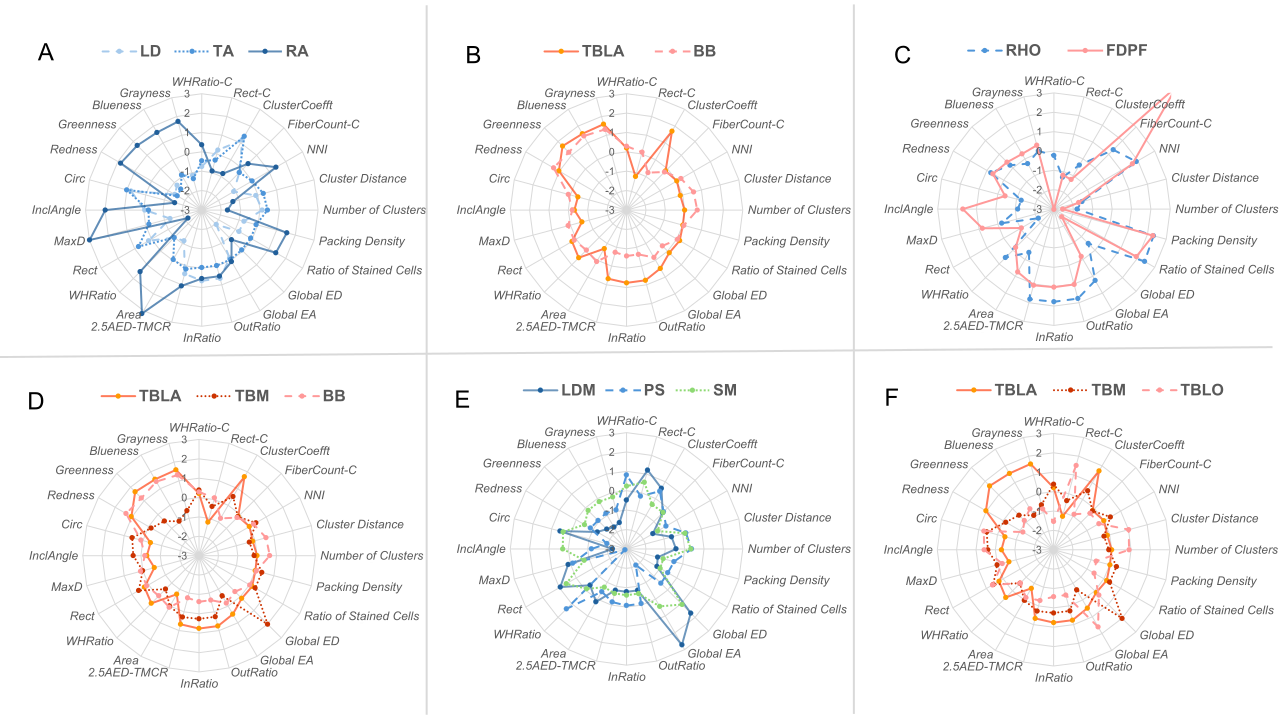
**Figure S12. Representative muscle comparisons reveal different spatial topology identities at similar type I fiber proportions.**

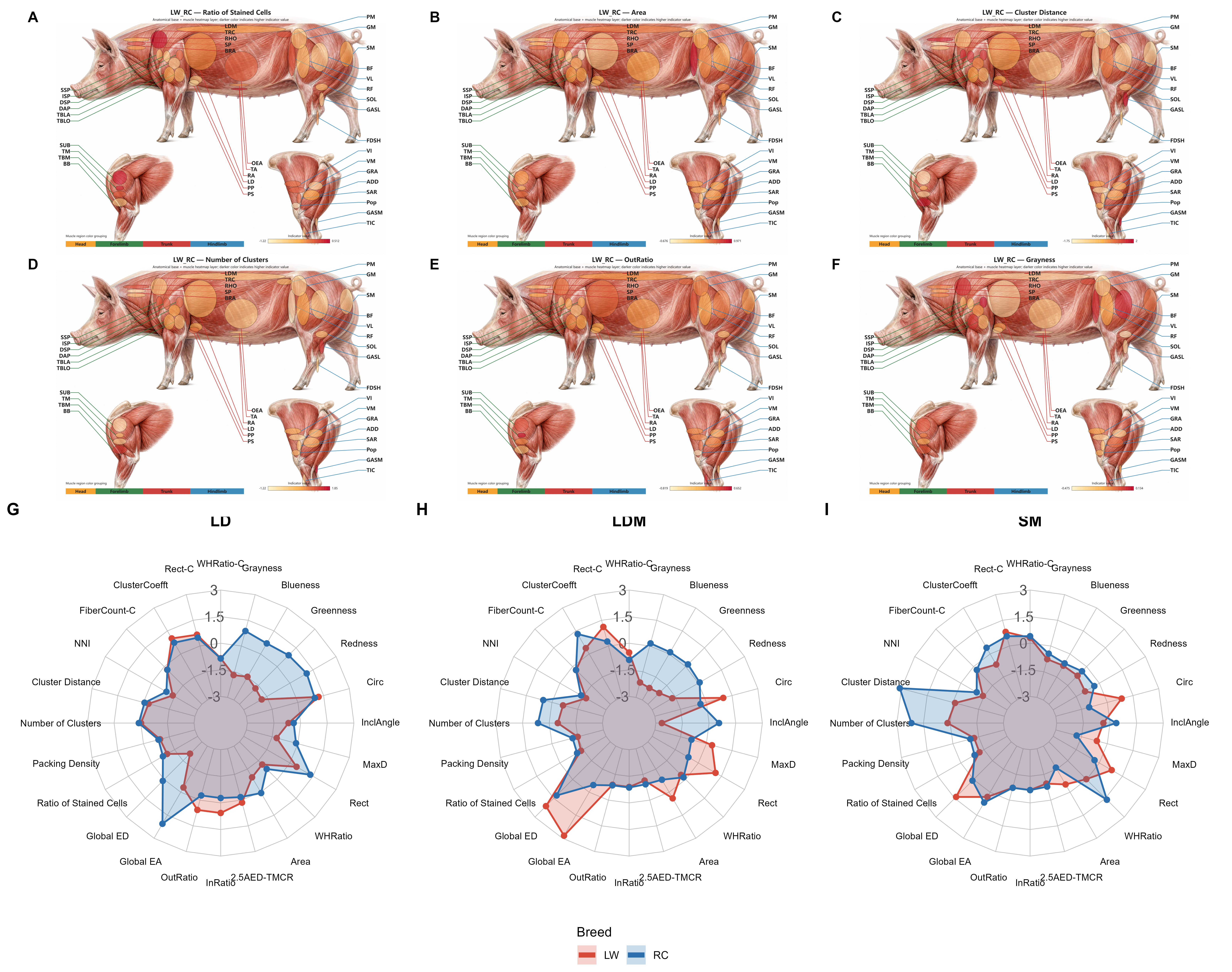
***Figure S13. Spatial phenotype differences across skeletal muscles in LW and RC pigs.*** *(A-F) The distribution of selected phenotypic indicators across all sampled skeletal muscles. (G-I) Phenotypic differences of specific skeletal muscles between two breeds of pigs.*

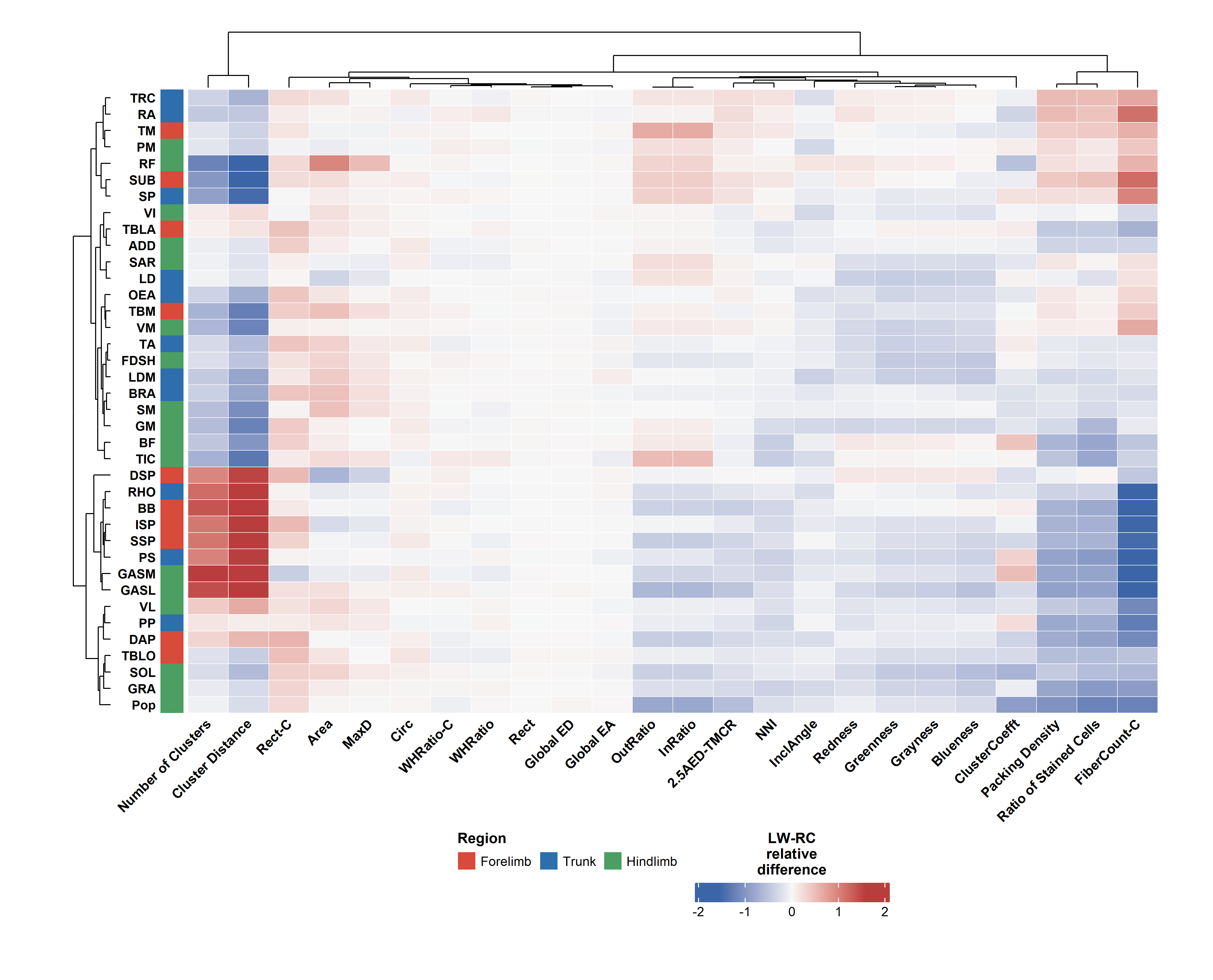
***Figure S14. Heatmap of breed-specific phenotypic differences across selected skeletal muscles.***

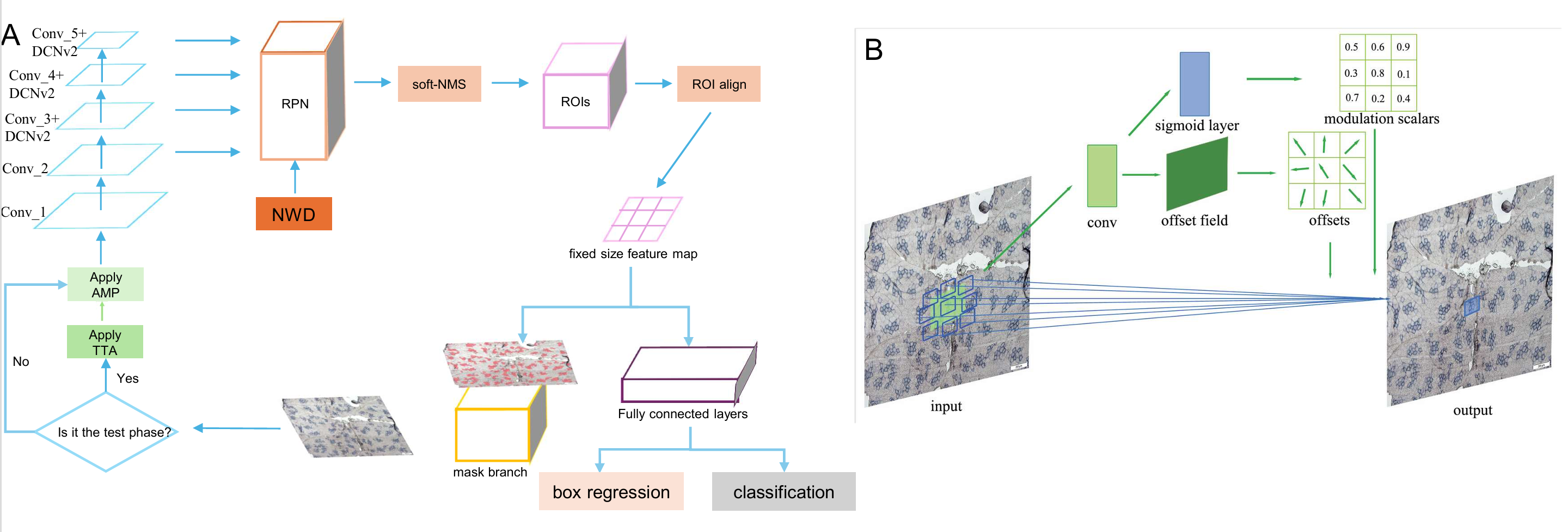

**Figure S15. Improved Mask R-CNN framework.**
